# Evidence that PV+ cells enhance temporal population codes but not stimulus-related timing in auditory cortex

**DOI:** 10.1101/213249

**Authors:** Bryan M. Krause, Caitlin A. Murphy, Daniel J. Uhlrich, Matthew I. Banks

## Abstract

Spatio-temporal cortical activity patterns relative to both peripheral input and local network activity carry information about stimulus identity and context. GABAergic interneurons are reported to regulate spiking at millisecond precision in response to sensory stimulation and during gamma oscillations; their role in regulating spike timing during induced network bursts is unclear. We investigated this issue in murine auditory thalamo-cortical (TC) brain slices, in which TC afferents induced network bursts similar to previous reports *in vivo*. Spike timing relative to TC afferent stimulation during bursts was poor in pyramidal cells and SOM+ interneurons. It was more precise in PV+ interneurons, consistent with their reported contribution to spiking precision in pyramidal cells. Optogenetic suppression of PV+ cells unexpectedly improved afferent-locked spike timing in pyramidal cells. In contrast, our evidence suggests that PV+ cells do regulate the spatio-temporal spike pattern of pyramidal cells during network bursts, whose organization is suited to ensemble coding of stimulus information. Simulations showed that suppressing PV+ cells reduces the capacity of pyramidal cell networks to produce discriminable spike patterns. By dissociating temporal precision with respect to a stimulus versus internal cortical activity, we identified a novel role for GABAergic cells in regulating information processing in cortical networks.

## Introduction

The timing of action potentials in cortical pyramidal cells often contains information about current and remembered sensory experiences (Abeles et al. 1994; Kayser et al. 2009; Lisman 2005; Nadasdy et al. 1999; Optican and Richmond 1987; Shmiel et al. 2005; Victor 2000; Wang et al. 2008). Spikes times in these codes may be referenced relative to temporal features of a stimulus, relative to an ongoing cortical oscillation, relative to other cells in a local ensemble, or a combination thereof. GABAergic interneurons play a role in regulating spike timing in pyramidal cells at the single cell level (Cobb et al. 1995; Pouille and Scanziani 2001) and at the level of homogeneous interneuronal networks (Whittington et al. 1995). It is unclear, however, how these observations translate to the diverse populations of cells that comprise even local cortical networks, i.e. the cortical microcircuit.

Four related issues contribute to this uncertainty. First, diverse groups of inhibitory cells regulate activity not only of pyramidal cells but also of each other. Thus, activation of excitatory afferents could produce net feedforward inhibition as at the single cell level (Pouille and Scanziani 2001), or net disinhibition, by activation of GABAergic interneurons that target other inhibitory cells (Pi et al. 2013; Zhang et al. 2016). For example, complex interactions between recurrent excitation and inhibition in local networks sharpen tuning in auditory cortex (Kato et al. 2017). Second, inhibitory influences on individual cells can differ from influences on an interconnected network due to nonlinearities present in the network (Seybold et al. 2015). Divisive inhibitory influences can become subtractive at the network level, and vice-versa. Third, excitatory inputs to the column, e.g. TC afferents, activate large numbers of cells simultaneously, triggering activity patterns that go beyond those observed in single cell recordings (Krause et al. 2014; MacLean et al. 2005). For example, although pyramidal cells in supragranular and infragranular layers both receive direct inputs from thalamus (Constantinople and Bruno 2013; Krause et al. 2014), differences in local excitatory and inhibitory connectivity and in integrative and firing properties position these two populations of pyramidal cells to play fundamentally different roles in signal coding and intrinsic cortical dynamics (Barth and Poulet 2012; Neske 2016; Sakata and Harris 2009).

Finally, activity within the cortical column is not a superposition of independent spike trains, but rather is highly correlated. Emerging evidence suggests that cortical information is conveyed via spatio-temporal patterns of spiking occurring within the context of coordinated network activity rather than by stochastic firing of individual cells (Abeles et al. 1993; Castejon and Nunez 2016; Luczak et al. 2015; Yuste 2015). Coordinated spiking indicative of bursts or packets of network activity with intervening periods of silence (ON/OFF periods; UP/DOWN states; synchronized state) was first described in cortex of sleeping and anesthetized animals (Neske 2016; Steriade et al. 1993). Arousal, especially in the form of locomotion or active sensation (e.g. whisking), is accompanied by a transition to the desynchronized state, corresponding to an extended ON period (McGinley et al. 2015; Poulet and Petersen 2008; Schneider et al. 2014; Zhou et al. 2014). Evidence suggests that the synchronized and desynchronized states represent points along a continuum (Curto et al. 2009; McGinley et al. 2015), and that the spatio-temporal patterns of spiking (‘packets’) observed during bursts in the synchronized state are preserved even during extended ON periods (Luczak et al. 2013). However, although the desynchronized state is associated with behavioral arousal, sensitivity is optimal when the network is otherwise quiescent, either in brief DOWN states or extended hyperpolarization (Curto et al. 2009; McGinley et al. 2015). Elevated firing rates when stimuli occur during optimal network states (Curto et al. 2009; Lakatos et al. 2005; McGinley et al. 2015) have been interpreted in terms of elevated burst probability when the network is in a DOWN state (Luczak et al. 2013).

Packets occur spontaneously in addition to being induced by afferent input, indicating that they can be powered by intracortical network dynamics. The importance of intracortical network mechanisms is evidenced by the resemblance between spontaneous bursts and activity triggered by sensory stimuli (Carrillo-Reid et al. 2015; Luczak et al. 2009; 2013; Miller et al. 2014; Sakata and Harris 2009). GABAergic cells play critical roles in regulating ongoing cortical network activity (Neske and Connors 2016), allowing its limited expression while preventing hyperexcitability (Destexhe 2010; Destexhe et al. 2003; Sanchez-Vives et al. 2010). Within this regulatory framework, what role do GABAergic cells play in constraining spike timing?

Multiple classes of cortical GABAergic cells have been identified in recent years, facilitating study of their roles in cortical processing (Tremblay et al. 2016). We focused on two of these classes, cells expressing parvalbumin (PV+), i.e. fast-spiking cells that target somata and proximal processes of pyramidal cells (Hu et al. 2014; Kawaguchi and Kubota 1998; Kubota et al. 2011), and cells expressing somatostatin (SOM+) that target distal dendrites of pyramidal cells (Kubota 2014; Markram et al. 2004). PV+ and SOM+ cells play distinct roles in regulating ascending and descending information streams in cortex. PV+ cells are strongly activated by TC afferents and constrain spike timing in pyramidal cells via rapid feedforward inhibition. SOM+ cells are postulated to modulate responses in distal dendrites to feedback excitation (Gentet et al. 2012). PV+ cells can limit spike output to narrow temporal windows by truncating EPSPs that would otherwise last for several milliseconds (Cruikshank et al. 2007; Gabernet et al. 2005; Pouille and Scanziani 2001). The same mechanism allows PV+ cells to control network synchrony and oscillatory behavior (Buzsáki and Wang 2012; Cardin et al. 2009; Sohal et al. 2009). Their strong activation during bursts in brain slices (Fanselow and Connors 2010; Neske et al. 2015; Tahvildari et al. 2012) suggests an important role in regulating intrinsic network activity. Here, we tested directly the role of SOM+ and PV+ cells in regulating network activity, and spike timing of pyramidal cells, using targeted recordings and optogenetics in auditory TC slices.

## Materials and Methods

### Mice and surgical procedures

All procedures were approved by the University of Wisconsin-Madison Animal Care and Use Committee and conform to American Physiological Society/National Institutes of Health guidelines. Mice were obtained directly or bred from stock (The Jackson Laboratory, Bar Harbor, ME). To identify specific types of interneurons, heterozygous SOM-tdTomato and PV-tdTomato mice were bred from homozygous Cre-dependent tdTomato (Stock 007914, Ai14) male and SOM-Cre (Stock 013044, SOM-IRES-Cre) or PV-Cre (Stock 008069, PV^cre^) female mice. Untargeted patch clamp recordings were also made from B6CBAF1/J mice (F1 hybrid of C57/B6 and CBA/J mice; The Jackson Laboratory). Some of the Cre-expressing animals were used for optogenetic activation/suppression experiments. In early experiments, this was accomplished via stereotaxic injection of adeno-associated virus expressing Cre-dependent halorhodopsin-YFP (AAV5/EF1α-DIO-eNpHR3.0-eYFP, Gene Therapy Center Vector Core, University of North Carolina at Chapel Hill, Chapel Hill, NC). Injections were performed on 3-5 week old mice of both sexes. Animals were anesthetized with isoflurane (1.5-2%) and craniotomized above auditory cortex based on stereotaxic coordinates (Franklin and Paxinos 2008). A total of 500-1000 nl of virus was injected into 2-3 sites spanning auditory cortex rostral-caudally approximately 500 μm from the lateral pial surface over 20-30 minutes (about 10 minutes per injection site). Injections were performed through a patch pipette broken to a tip diameter of approximately 50 μm and controlled with a Nanoject II (Drummond Scientific Company, Broomall, PA) mounted on a stereotaxic frame. Injected mice recovered for 3-5 weeks prior to preparation of brain slices. In later experiments, we bred SOM-Cre and PV-Cre animals with mice with Cre-dependent expression of the inhibitory pump archaerhodopsin (Stock 021188, Ai40(RCL-ArchT/EGFP)-D) to yield pups with ArchT expressed selectively in each interneuron subtype.

### Slice preparation

Auditory TC slices were prepared from male or female mice (*de novo*: 4-12 week old; recovered from surgery: 6-10 week old) deeply anesthetized with isoflurane and decapitated. We modified a previously described slicing method (Cruikshank et al. 2002; Krause et al. 2014) to maximize preservation of afferents connecting auditory thalamus and cortex in young adult and adult animals. In this method, brains were blocked with three cuts (Supplementary Figure 1), each perpendicular to the preceding cut: first, a sagittal cut about 2 mm lateral to the midline, preserving the larger block; second, a 45° rostral-dorsal to caudal-ventral cut, preserving the dorsal/caudal block; and third, a cut 20° off the horizontal-caudal plane, preserving the lateral/caudal block. The resulting block was affixed to a vibrating microtome on the final cut face and sliced into 500 μm sections. Slices at this angle are roughly midway between the coronal-horizontal plane, such that ‘caudal’ regions of the slice are about equally ‘dorsal,’ but tilted slightly so that medial regions are ventral to lateral regions. In the adult mouse, this blocking procedure best preserves the ventral medial geniculate nucleus (MGv) and primary auditory cortex (Au1) in the same slice along with the C-shaped TC fibers that travel rostal and ventral before arcing caudal on the way to auditory cortex (based on the Allen Mouse Brain Connectivity Atlas, http://connectivity.brain-map.org)(Oh et al. 2014).

During blocking and sectioning, slices were maintained in ice-cold cutting ACSF consisting of (in mM) 111 NaCl, 35 NaHCO_3_, 20 HEPES, 1.8 KCl, 1.05 CaCl_2_, 2.8 MgSO_4_, 1.2 KH_2_PO_4_, 10 glucose and bubbled with 95% O_2_/5% CO_2_. Once cut, slices were immediately placed in cutting ACSF warmed to 34°C, which cooled to room temperature as slices rested for at least one hour. Slices were moved to the recording chamber and perfused at >6 ml/min with regular ACSF consisting of (in mM) 111 NaCl, 35 NaHCO_3_, 20 HEPES, 1.8 KCl, 2.1 CaCl_2_, 1.4 MgSO_4_, 1.2 KH_2_PO_4_, and 10 glucose, warmed to 30-33°C and bubbled with 95% O_2_/5% CO_2_. Maintaining a strong flow of ACSF over the slice by minimizing the volume in the chamber and strategic arrangement of inflow and outflow was necessary to ensure robust network activity.

### Electrophysiology and data analysis

Bipolar stimulating electrodes (100KΩ, FHC) were placed into the TC fiber bundle rostral to hippocampus; we have shown previously that this stimulation configuration activates current sinks in auditory cortex indistinguishable from those elicited by stimulation in thalamus (Krause et al. 2014). Current pulses consisted of a 200 μs biphasic square wave of amplitude 10-100 μA (STG4002 stimulator, Multichannel Systems, Reutlingen, Germany). The vertical strip of cortex (‘column’) with optimal responses to the stimulated TC fibers was identified by recording TC responses in layer 4 of Au1 at 250 μm increments, using a glass patch pipette filled with ACSF and broken to a resistance of 500-700 kΩ. We used these responses to identify the cortical location with the largest early (<10 ms latency) extracellular responses; further recordings were focused at this location. During data collection, the stimuli were delivered as a train of 4 pulses at 40 Hz, which reliably evoked recurrent network activity (Krause et al. 2014). The stimulus intensity used for a given experiment was adjusted to give network bursts that occurred reliably after pulse 2 and before pulse 4 in the train; the specific intensity required likely depended on the integrity of TC fibers in each slice. For most experiments, there was substantial adaptation after the first or second trial so we omitted the first two trials. To further account for adaptation of responses, we repeated our analyses using only the latest half of the trials, but this did not qualitatively impact results or conclusions.

For single-cell recordings, SOM+ or PV+ interneurons were patched using a combination of fluorescence and differential interference contrast (DIC) microscopy. Light from a mercury arc lamp (X-Cite exacte; Lumen Dynamics, Mississauga, Ontario, Canada) passed through an excitation filter (540-580 nm; Chroma, Bellows Falls, VT) and broad spectrum transmitted light were presented simultaneously to identify labeled cells and surrounding tissue features, respectively, and emission/transmittance (emission filter 593-667 nm; Chroma) captured on a CCD camera (C9100-02, Hamamatsu Corp., Sewickley, PA). A borosilicate micropipette (KG-33, 1.7 mm OD, 1.1 mm ID; King Precision Glass, Claremont, CA), pulled to give open-tip resistance of 3-5 MΩ (P-1000; Sutter Instruments, Novato, CA) filled with intracellular solution (in mM: 140 K-gluconate, 10 NaCl, 10 HEPES, 0.1 EGTA, 2 MgATP, and 0.3% biocytin; pH 7.2), was advanced to contact the targeted cell and a >1GΩ seal made with weak negative pressure. Pyramidal cells were patched similarly, with transmitted light only, and identified based on their visible triangular morphology with apical dendrite under DIC optics. Spikes were recorded in the on-cell configuration in voltage-clamp in response to trains of 4x40 Hz TC stimuli. After on-cell recording, whole-cell access was established and recordings continued in current-clamp mode. For some cells, whole-cell access occurred before on-cell recordings were completed. In these cells, spikes recorded in whole-cell configuration were analyzed. In addition to Cre-dependent tdTomato expression and morphological characteristics, cell types were identified by their characteristic responses to current pulses, particularly spiking patterns and rates (i.e., regular-spiking vs. fast spiking). For all recordings, the recorded voltage or current signal was low-pass filtered at 4 kHz and digitized at 40 kHz.

During all experiments, including during single-cell recordings, simultaneous extracellular population recordings in layer 5 (and in some experiments also in layers 2/3) were used to measure network activity. Broken glass pipettes (as above, broken to resistances of 500kΩ-800kΩ) filled with regular ACSF were inserted into layer 5 at a cortical depth slightly greater than halfway from pia to white matter. Stimulus artifacts were blanked by interpolating between points before and after the artifact. Extracellular voltage was filtered between 500-3000 Hz, full-wave rectified, and smoothed by convolution with a Gaussian kernel with unit integral and σ=2 ms to produce a smoothed MUA signal (smMUA), also referred to as “population activity” throughout this paper. A threshold for elevated activity was defined as the geometric mean of all the points greater than the arithmetic mean of the smMUA signal (Krause et al. 2014; Sakata and Harris 2009). Burst onsets were defined as periods above threshold for at least 80% of points in a 20 ms window, and burst offsets defined as a decrease below the threshold for 80% of points in a 50 ms window. These criteria reliably identified network bursts observed by eye, and did not include early responses to individual stimulus pulses (which are too brief).

Many measures have been suggested for quantifying spike timing, but creating a firing-rate independent measure of correlative features such as spike timing across trials is nontrivial (Cohen and Kohn 2011; Cutts and Eglen 2014; Joris et al. 2006). We assayed spike timing using a variation of the previously-described “spike-time tiling coefficient” (STTC; (Cutts and Eglen 2014)), which computes the fraction of spikes coincident with a specified coincidence window (Dt) above those expected by chance. Instead of comparing between two simultaneously recorded units as originally described, the measure was computed between all permutations of trials for a given unit. The calculation depends on a free parameter Δt, which defines the window on which spikes are considered “coincident.” Then, two proportions are calculated, P_A_, the proportion of spikes from spike train A that are within Δt of any spike in spike train B, and T_B_, the proportion of all time that is within Δt of any spike in spike train B. The STTC is calculated as:

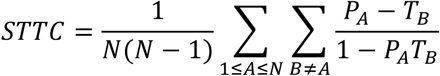

This measure is high (approaching 1) when all spikes occur at the same time as spikes in other trials, and zero for randomly distributed spikes. The STTC was only calculated for cells that fired at least one spike on more than 1/3 of trials. The STTC tends toward unity as the coincidence interval increases. Importantly, the STTC is robust to changes in firing rate (Cutts and Eglen 2014), but will change with coordinated changes in firing rate that are on time scales much slower than Δt.

A second measure of spiking precision across trials was adapted from a spike train distance metric (Victor and Purpura 1996). We define the spike train similarity S between two trials (*a,b*) as:

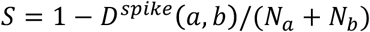

where *D*^*spike*^ is the Victor-Purpura metric for spike train distance (Victor and Purpura 1996), defined as the cost of transforming one spike train into another by adding, shifting, or removing spikes. The cost of adding or removing a spike is defined as 1; the cost of shifting a spike is equal to *q*·Δt, where *q* is a free parameter that determines the temporal precision of interest. We used *q*=0.5/ms to compare with our STTC results, meaning that it is more efficient to remove and add a spike (a total cost=2) than to shift a spike by more than 4 ms. We normalized the Victor-Purpura distance by the sum of the number of spikes *N* in trains *a* and *b* (Dimitrov et al. 2014), and subtracted from 1 to transform from a measure of distance to a measure of similarity. We defined the spike train similarity for one cell across all trials as the average of *S* across all trial combinations (*a,b*) for which *N*_*a*_+*N*_*b*_≠*0*. Even with the normalization to number of spikes, this measure can be sensitive to changes in firing rates, but has an advantage over the STTC measure in that it uses a non-binary measure of coincidence.

We wished to compare across experiments the timing of spikes relative to burst onset and offset. Because population bursts vary in duration between and within slices, we normalized burst duration and defined the “burst phase” as 0 at the start of a burst and 1 at the end of a burst. The “burst firing phase” was the fractional time of each spike between onset and offset of the burst detected on that trial. To calculate firing rates during bursts, spike trains recorded from single cells were convolved with a Gaussian kernel (σ=2 ms) before temporal scaling to units of burst phase to preserve units of firing rate in terms of spikes/second (Hz).

Statistical comparisons between cell types used standard one-way ANOVA when normality was not rejected using a one-way Kolmogorov-Smirnov test; otherwise, a Kruskal-Wallis test of analysis of variance by ranks was used. Levene’s test was used to compare variances. All statistical tests used the MATLAB (Mathworks, Natick, MA) Statistics Toolbox.

### Optogenetic suppression of inhibition

Halorhodopsin or ArchT pumps were activated by passing light from the arc lamp through the microscope objective (10x) centered on the cortical region of study using a filter in the excitation range for halorhodopsin (540-580 nm). Activation of these constructs caused substantial hyperpolarization in labeled cells (see Supplementary Figure 3). The light was turned on 100 ms before stimulus onset and held on for a total of 500 ms. Light intensity was titrated for each experiment to a level that produced an effect on population responses, in the range of 0.6 to 2.9 mW/mm^2^ for halorhodopsin and 1.5 to 5.8 mW/mm^2^ for ArchT. Light-on trials were interleaved with light-off trials. The first two trials of each type were discarded because these trials often contained induced bursts that differed substantially in magnitude and latency from subsequent events. The effects of optogenetic suppression on individual cells were tested with Wilcoxon signed-rank tests.

We did not observe any evidence of toxicity, for example abnormal “blebbing” of membranes, that has been associated with older halorhodopsin constructs (Gradinaru et al. 2008). Additionally, patched eNpHR3.0+ cells had normal resting potentials and firing behavior. Our stimuli were brief (several hundred milliseconds) relative to durations known to impact intracellular chloride concentrations which span several seconds (Raimondo et al. 2012). We also did not observe any differences between eNpHR3.0 and ArchT slices with or without light stimulation, and did not observe any differences without light in those slices compared to slices without any optogenetic expression.

### Linear mixed-effects model

To analyze the effect of optogenetic suppression of different interneuron populations, we fit population responses using a linear mixed-effects model (Kristensen and Hansen 2004; Winter 2013). This approach allows for controlling for random effects due to repeated measures within subjects, enabling us to consider non-independent samples in a principled way. Other methods such as using repeated-measures ANOVA could not be structured to our data without enforcing independence by summarizing each experiment with a single value for each condition (such as a mean or median) and therefore losing important information about within-subject variability or consistency.

Models consisted of a response variable based on the population activity (smMUA peak, burst latency, or burst duration), fixed effect of interneuron suppression (LightOff/SOM+/PV+) and recording layer (L5 or L2/3), an interaction term between the two fixed effects, and random effect of slice (experiment), with random slopes by condition and by layer for the effect of slice. In summary, the full model was as follows, with fixed effects β, random effects γ, suppression *i,* layer *j,* and slice *k*:

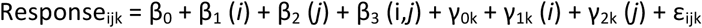

Statistical evaluation of the linear mixed effects model occurred at two levels. First, we used a likelihood ratio test to determine whether fixed effects in the above equation are significant, comparing the full model to a reduced model that lacks the interaction term (the interaction term indicates an effect of suppression that depends on layer). If omitting the reaction term significantly reduced model likelihood, we further analyzed the individual coefficients for that term. Otherwise, we omitted the interaction from the model. For example, for the effect of suppression of interneurons on the peak and latency (Figure 6; Table 2), the model was significantly improved by including the interaction between interneuron suppression and layer (i.e., β_3_; peak χ^2^(2)=124.1, p<0.0001; latency χ^2^(2)=31.7, p<0.0001). For the duration, the interaction was not significant (χ^2^(2)=0.72, p=0.69) but both of the individual fixed effects (β_1_, χ^2^(2)=20.2, p<0.0001, and β_2_, χ^2^(2)=17.4, p<0.0001) were significant. This result indicates that, although there were effects of interneuron suppression and differences between layers, there was no difference in the effect of interneuron suppression across layers. Therefore, we used a reduced model with no interaction term for evaluating duration and used the full model for the other measures. We report likelihood ratio tests using chi-squared values. Residuals were visually inspected to confirm homoscedasticity. For the latency and duration measures, heteroscedasticity was corrected by log-transforming the response variables. After choosing the appropriate models, we tested the significance of individual coefficients (Table 2). For ease of interpretation, coefficient estimates for these models were exponentiated after fitting to express effects as multiplicative gains. Coefficients are reported with 95% confidence intervals. All data analysis and statistical comparisons used the MATLAB (Mathworks, Natick, MA) Statistics Toolbox and custom MATLAB software.

**Figure 6.**
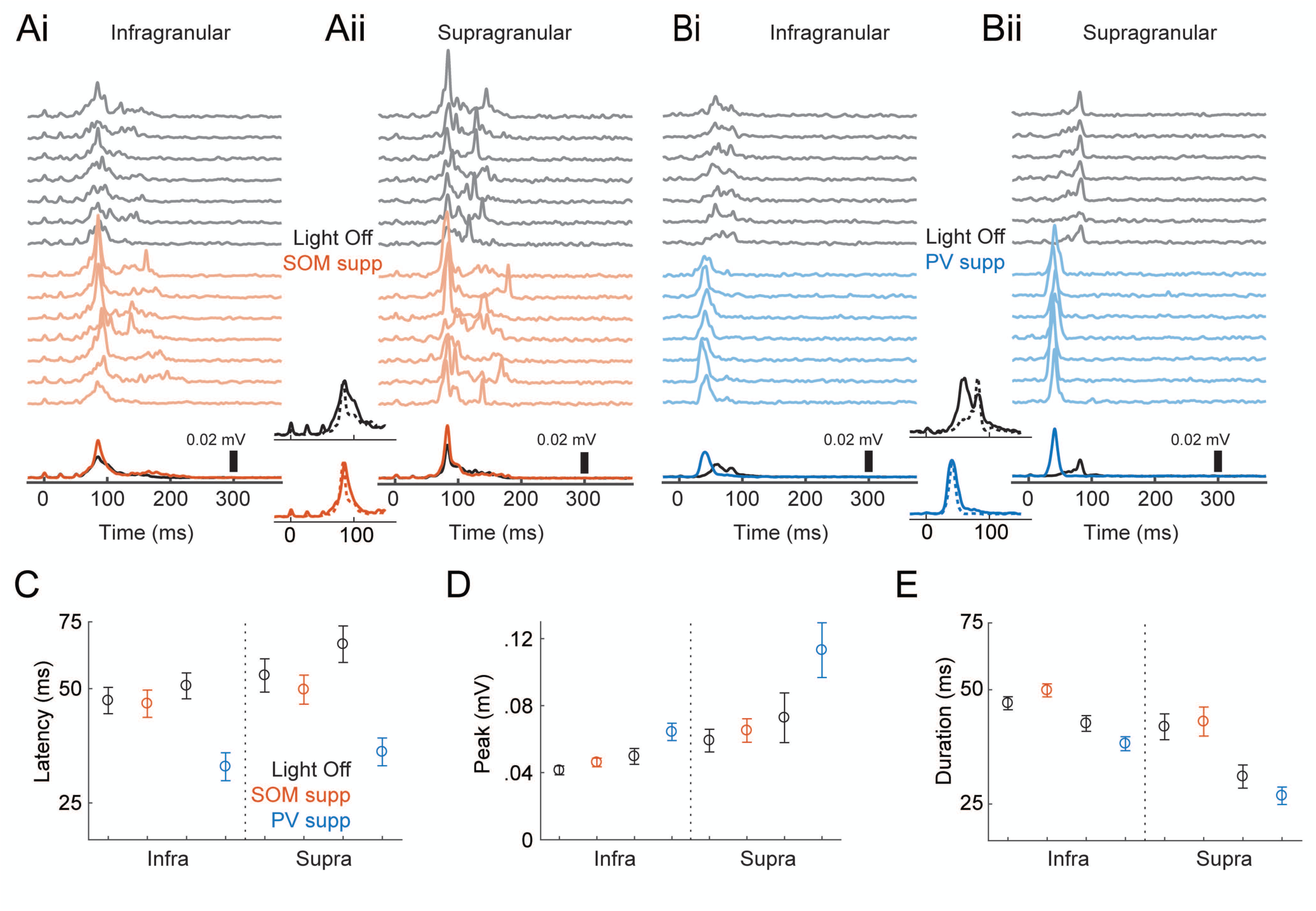
Effects of optogenetic suppression of interneurons on network bursts. (Ai) Population activity (smoothed MUA) recorded in layer 5 in an animal expressing an inhibitory opsin in SOM+ cells. The top gray traces are single trial examples recorded with no light; the colored traces are with yellow light suppressing SOM+ cells. Darker black/colored lines at the bottom are means across trials. Light came on 100 ms before activation of TC afferents (TC afferents were activated starting at time=0) and lasted a total of 500 ms. (Aii) Simultaneously recorded data from supragranular layers 2/3 in the same slice as in (A). Central insets show the same data, but grouped by control (black, top) and SOM+ suppression (colored, bottom) and normalized by peak to emphasize temporal relationships, with infragranular bursts as solid lines and supragranular bursts as dotted lines. (Bi-ii) Another example experiment, but with PV+ suppression. Conventions are the same as in (Ai-ii). (C-E) Means ± standard error for peak (C), latency (D), and duration (E). Results are separated according to the layer (infragranular/supragranular) where population activity was recorded; within each layer, the black, red, and blue symbols represent light off, SOM+ suppressed, and PV+ suppressed, respectively.

### Tempotron learning model

To test how discriminable random spiking patterns are with differing distributions of burst firing phase, we implemented a tempotron learning model (Gütig and Sompolinsky 2006). Briefly, the tempotron is a single-compartment, leaky integrate-and-fire neuron that responds in a binary fashion (“spike” or “no-spike”) to a pattern of weighted synaptic inputs. Patterns were arbitrarily assigned to “go” or “no-go” categories, where the correct response to a “go” pattern is a spike, and the correct response to a “no-go” pattern is no spike. The membrane voltage V(t) was given by summing the weighted inputs convolved with a causal filter K given by the equation:

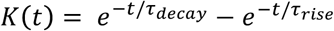

We set *τ*_*decay*_ := 15 *ms* and *τ*_*rise*_ := 3.75 *ms*. If V(t) reached threshold, the trial ended at time t and a spike was registered. After each trial, weights ω were updated according to the tempotron learning rule:

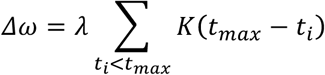

This rule multiplies the maximum update step (λ := 0.001 ∗ ∫ k) by a factor that represents the contribution of spikes at the observed times relative to the time of the maximum V(t). Synaptic weights were increased by ∆ω when there was a miss error and decreased by ∆ω when there was a false alarm error. We also included a momentum µ := 0. 99, as in the original tempotron model, to accelerate the learning process. Because deep cells had higher mean firing rates than superficial cells, in some simulations (see Results) we matched firing rates between deep and superficial cells by omitting deep cells in descending rank order until the mean firing rate in the sample of deep cells was as close as possible to the mean firing rate of the superficial cells.

## Results

### Induced network activity

We seek to understand the spatio-temporal patterns of spiking during correlated network activity (‘network bursts’) within the cortical column, and the regulation of this activity by two classes of cortical inhibitory cells. We chose to study these questions in brain slices (Figure 1A), in which we and others have shown that such activity is readily observed. In murine auditory TC slices, two types of responses are observed in response to activation of TC afferents. Short latency current sinks in layer 4 and EPSPs in cells of all recorded layers are consistent with monosynaptic TC inputs (Cruikshank et al. 2002; Krause et al. 2014; Raz et al. 2014) (Figure 1B, inset). Longer latency, variable duration responses, corresponding to network bursts, are also observed. Intracellularly, these consist of depolarizations that arise from polysynaptic inputs and are shared across cells in the column; spiking activity recorded in individual cells and as multiunit activity extracellularly occurs preferentially during these burst events (Figure 1B). Here, we defined network bursts as a sustained increase in the smoothed population MUA recorded in layer 5 (Figure 1C; see Methods). Burst duration varied with stimulus intensity and across slices, but was typically around 40-70ms (Figure 1D). Network bursts of similar duration have been observed previously *in vitro* (Hentschke et al. 2017; Krause et al. 2014; MacLean et al. 2005; Metherate and Cruikshank 1999; Shu et al. 2003) and in auditory cortex in response to acoustic stimuli *in vivo* (Curto et al. 2009; Luczak et al. 2009; 2013; Sakata and Harris 2009).

**Figure 1.**
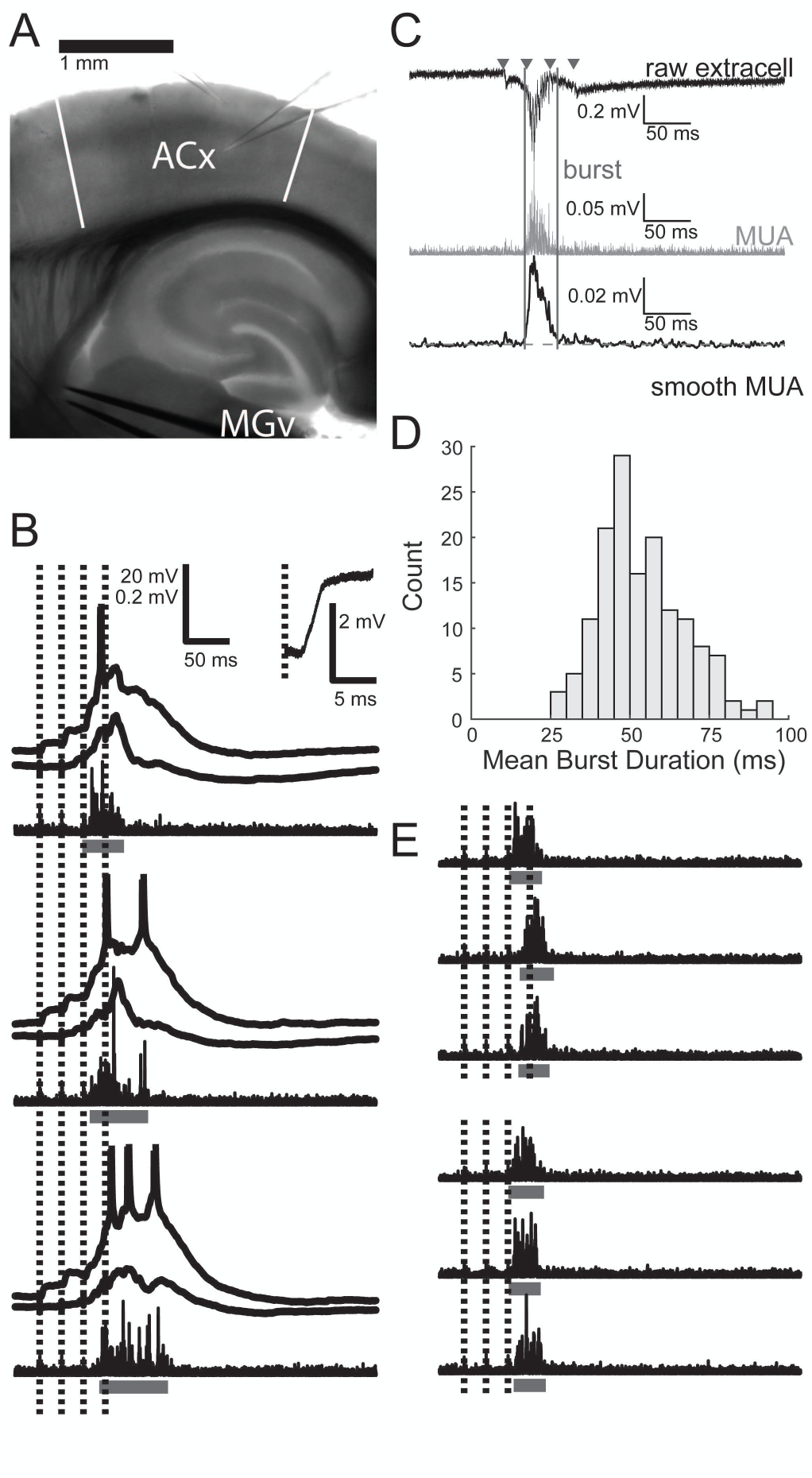
Induced network bursts and spiking activity. (A) Typical recording configuration. Two glass micropipettes are shown, one in layer 2/3 and one in layer 5. During single-cell recordings, another electrode always monitored population activity in layer 5. A metal stimulating electrode pair is placed in the fiber tract from MGv to auditory cortex. Toward the left of the image is approximately rostral/ventral; toward the top is roughly lateral, but note from Supplementary Figure 1 that the slicing procedure results in slices not in any cardinal plane of section. ACx: auditory cortex, with boundaries indicated by white bars; MGv: medial geniculate nucleus, ventral division. (B) Three example trials from two simultaneously recorded cells, along with high-pass filtered population activity recorded on a third electrode. Horizontal bars indicate bursts detected in extracellular activity. Inset shows magnified response to the first stimulus pulse from one cell. EPSP latency = 2.4 ms. (C) Network burst detection procedure. The raw extracellular trace (black, top) is filtered between 500-3000 Hz to give the MUA signal (grey, center). TC stimulus times are marked with triangles. The MUA signal is smoothed with a Gaussian kernel to give the smoothed MUA signal (black, bottom). Burst onset and offset (vertical lines) are determined by threshold (horizontal dashed line) crossings above the geometric mean of the smoothed MUA signal (see Methods). (D) Histogram of mean burst durations while recording from each individual cell. (E) Six example trials and detected bursts. On three trials, the fourth stimulus pulse (which would have occurred during bursts) is omitted, but there is no change in burst duration. Scale same as in (B).

Afferent stimulation can itself induce bursts of correlated network activity within the auditory cortical column. Importantly, these stimuli can also coincide with spontaneous bursts, or with bursts induced by prior stimulation (Destexhe and Pare 1999; Luczak et al. 2013; MacLean et al. 2005; Petersen et al. 2003; Rigas and Castro-Alamancos 2009). Our observation that nearly all spiking activity occurs in the context of network bursts suggests that driving the network into the activated state underlying network bursts is necessary for throughput of afferent information. Thus, we are interested in how the network responds to afferent stimulation occurring during bursts. To investigate this, we chose for most of our experiments trains of afferent stimuli (4x40 Hz, 10-100µA) at intensities that induced bursts after the second and prior to the fourth stimulus in a train of four pulses, such that there was always at least one stimulus pulse occurring during the burst (Figure 1B).

Because stimulation during ongoing bursts can hasten their termination, we verified that the bursts we observe are not short merely due to our experimental paradigm that includes trains of stimuli that overlap with induced bursts. We found no difference in burst duration with versus without stimulation amidst bursts in a subset of experiments (n=7, mean duration difference ± SEM= 1.1±5.0 ms; paired t-test p=0.84; Figure 1E). Duration of bursts can also be affected by the method for detecting and defining bursts, which differs across studies. For example, the duration of the thresholded MUA signal in layer 5 was typically less than the intracellular depolarization signal (Figure 1B), though we note that the vast majority of spiking activity occurred within the detected bursts according to our definition (Hentschke et al. 2017; Krause et al. 2014).

### Cell type-specific firing patterns during network bursts

To investigate the role of GABAergic cells in regulating network activity, and specifically spike timing in the context of this activity, we first determined the firing patterns of pyramidal, PV+, and SOM+ cells during network bursts induced by thalamic stimulation. To record from specific classes of GABAergic cells during network bursts, we prepared slices from transgenic animals expressing the fluorophore tdTomato in either PV+ or SOM+ cells (Supplementary Figure 2). We identified pyramidal cells based on their morphology in these same slices. The three cell types had distinct patterns of firing with respect to bursts (Figure 2A-C) and intrinsic properties (Table 1; see also Supplementary Material). Pyramidal cells tended to fire sparsely during bursts (Figure 2A). By contrast, GABAergic cells tended to fire more densely. SOM+ cells tended to fire multiple spikes late in bursts; occasionally, some SOM+ cell spikes occurred after the detected burst duration (Figure 2B; 13.4% of SOM+ cell spikes were after bursts, compared to only 2.6% of pyramidal cell spikes and 3.9% of PV+ cell spikes). PV+ cells tended to fire multiple spikes early and throughout bursts (Figure 2C). On average, pyramidal cells fired significantly fewer spikes per trial (including spikes before or after detected bursts) than either inhibitory cell population (Figure 2D; H(2)=32.1, p<0.0001; Pyr vs. SOM+ p<0.0001, Pyr vs. PV+ p<0.0001) and were more likely to fire no spikes on a given trial (Figure 2E; H(2)=21.7, p<0.0001; Pyr vs. SOM+ p=0.0003, Pyr vs. PV+ p=0.0016). Thus, although interneurons make up only 10 – 20% of neurons in auditory cortex, their substantial firing activity positions them to exert strong influence over induced network activity. Pyramidal cells in layer 5 fired more spikes per trial than pyramidal cells in layers 2/3 or 4 (not shown; H(2)=21.0, p<0.0001; Pyr L2/3 vs. L5 medians 0.06 vs. 1.0, p=0.0008; Pyr L4 vs. L5 medians 0 vs 1.0, p=0.0009). There were no significant laminar differences in firing rate for either interneuron type.

**Figure 2.**
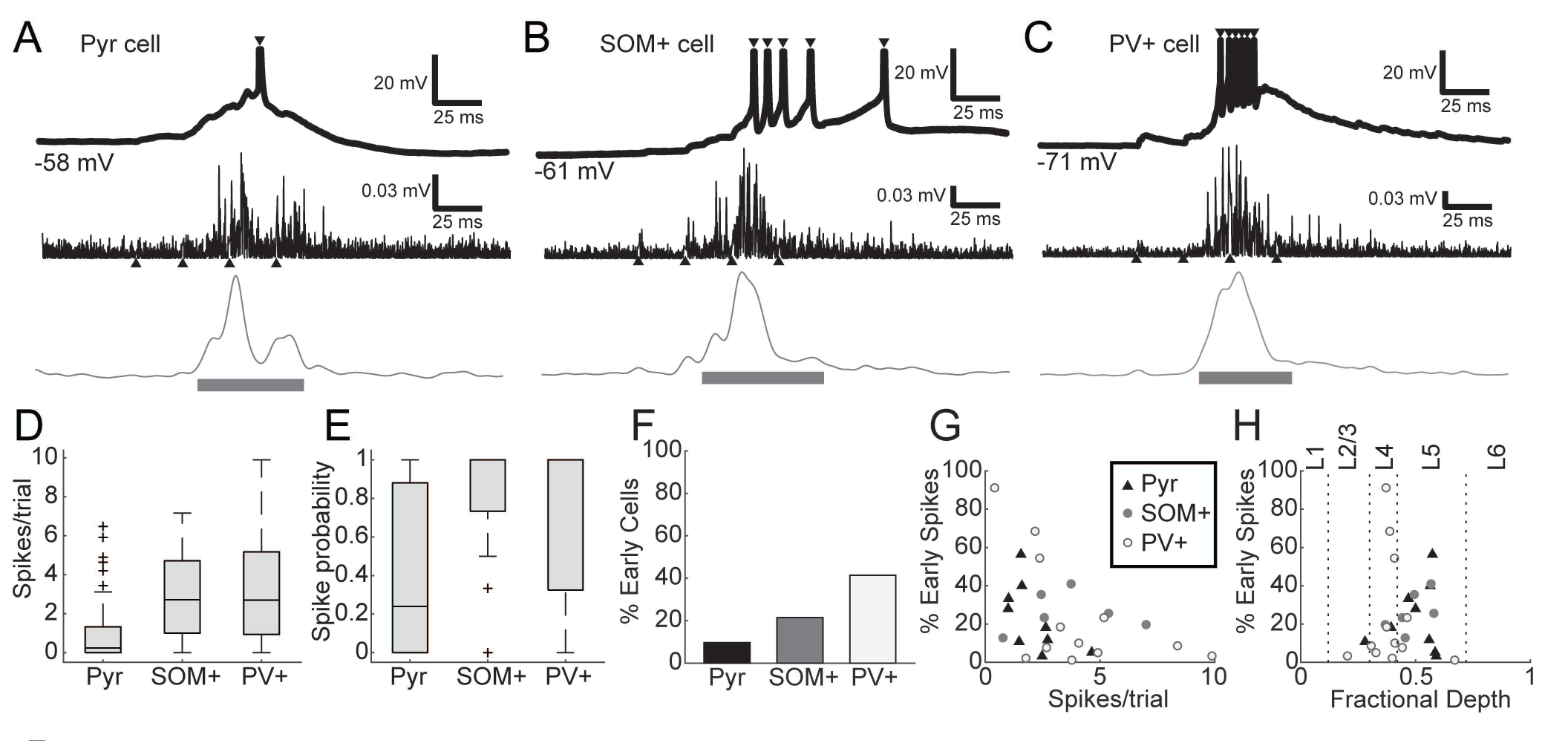
(A-C) Intracellular recordings (top) from a pyramidal (A), SOM+ (B), and PV+ cell (C), along with example MUA traces (middle: high-pass filtered and rectified signal; bottom: smoothed MUA signal) from three different slices . Upward-pointing triangles mark the time of TC afferent stimulus pulses, and downward pointing triangles mark intracellular spikes. Horizontal bars indicate duration of detected bursts. Spikes are truncated above −20 mV. Values in mV indicate resting membrane potential. (D) Pyramidal cells averaged fewer spikes per trial than SOM+ or PV+ cells. Box plots here and elsewhere indicate median (horizontal line), interquartile range (box), range outliers (black bars), and outliers (symbols). (E) Most pyramidal cells (Pyr) fired sparsely or not at all, even during UP states, whereas most interneurons (SOM+ and PV+) fired at least one spike on every trial. (F) A minority of all cell types fired some fraction of their spikes early, prior to burst onset, though this was more common in interneurons, particularly PV+ cells. (G) Even among those cells that fired some early spikes, most cells fired most of their spikes during induced bursts. (H) PV+ cells with the greatest fraction of early spikes were found in layer 4; pyramidal and SOM+ cells that fired early spikes were mostly found in layer 5. Horizontal axis depicts fractional depth from pia to white matter; dotted lines mark boundaries between layers 1, 2/3, 4, 5, and 6.

**Table 1.** Properties of recorded cell types. Values are presented as mean±SD. Spike widths are full-width at half-maximum. ISI ratio is a measure of spike adaptation equal to the ratio of first to last interspike interval for a 400 ms depolarizing current at half-maximal firing rate. Resting membrane potential (RMP) did not vary significantly between groups (F(2,112)=1.86, p=0.16). Input resistance was significantly higher in SOM+ cells (F(2,112)=21.0, p<0.0001; Pyr vs. SOM+, p<0.0001; Pyr vs. PV+, p=0.69; SOM+ vs. PV+, p<0.0001). Membrane time constants were significantly shorter in PV+ cells (F(2,112)=14.0, p<0.0001; Pyr vs. SOM+, p=0.10; Pyr vs. PV+, p=0.0002; SOM+ vs PV+, p<0.0001). Spike widths were significantly different between all groups (F(2,112)=84.3, p<0.0001; Pyr vs. SOM+, p<0.0001; Pyr vs. PV+, p<0.0001; SOM+ vs PV+, p<0.0001). Spike adaptation (ISI ratio) was significantly different between all groups (F(2,112)=89.6, p<0.0001; Pyr vs. SOM+, p<0.0001; Pyr vs. PV+, p<0.0001; SOM+ vs PV+, p=0.0001). EPSP slopes were greater in PV+ cells (H(2)=30.6, p<0.0001; Pyr vs. SOM+, p=0.69; Pyr vs. PV+, p<0.0001; SOM+ vs PV+, p<0.0001). Similarly, EPSP latencies were shorter in PV+ cells than pyramidal or SOM+ cells (H(2)=20.2, p<0.0001; Pyr vs. SOM+, p=0.75; Pyr vs. PV+, p=0.0002; SOM+ vs PV+, p=0.0003).

| | RMP (mV) | $R_{in}$ (MΩ) | $\tau_m$ (ms) | Spike width (ms) | ISI ratio | EPSP slope (mV/ms) | EPSP Latency (ms) |
| --- | --- | --- | --- | --- | --- | --- | --- |
| Pyr | -69.1±10.5 | 92.6±53.2 | 10.2±6.2 | 1.04±0.26 | 0.30±0.18 | 2.35±2.39 | 2.6±0.86 |
| SOM+ | -64.6±5.2 | 173.9±84.5 | 13.4±9.6 | 0.67±0.25 | 0.61±0.26 | 3.99±7.86 | 3.4±2.4 |
| PV+ | -66.8±11.4 | 82.0±26.0 | 4.2±1.3 | 0.37±0.14 | 0.86±0.15 | 15.0±12.2 | 1.9±0.36 |

**Table 2.** Population effects of optogenetic suppression of interneurons during UP states. Intercepts reflect population responses recorded in L5 with no light. Interpretations indicate the meaning of a given parameter based on the ordering of parameters in the model; β-parameters not listed are those defined as zero. Units for the peak measure are in mV. For burst latency and duration measures, the response variables were log-transformed prior to regression; estimated coefficients are reported in their exponentiated form (see Methods) and represent unitless multiplicative factors; the intercepts are expressed in units of ms.

| Measure/Parameter | Interpretation | Estimate | 95% CI | p-value |
| --- | --- | --- | --- | --- |
| LATENCY | *Intercept* | 48.4 | [43.0 54.5] |  |
| $\beta_1(\text{SOM}_{\text{supp}})$ | Effect in L5 of SOM+ suppression | 0.96 | [0.93 0.99] | 0.014 |
| $\beta_1(\text{PV}_{\text{supp}})$ | Effect in L5 of PV+ suppression | 0.62 | [0.56 0.68] | <0.0001 |
| $\beta_2(\text{L2/3})$ | L2/3 vs. L5, with no light | 1.15 | [1.09 1.21] | <0.0001 |
| $\beta_3(\text{SOM}_{\text{supp}}, \text{L2/3})$ | Interaction between SOM+ suppression and L2/3 recording | 0.95 | [0.93 0.97] | <0.0001 |
| $\beta_3(\text{PV}_{\text{supp}}, \text{L2/3})$ | Interaction between PV+ suppression and L2/3 recording | 0.94 | [0.90 0.97] | 0.0003 |
| PEAK | *Intercept* | 0.0442 | [0.0388 0.0497] |  |
| $\beta_1(\text{SOM}_{\text{supp}})$ | Effect in L5 of SOM+ suppression | 0.0041 | [0.0009 0.0072] | 0.011 |
| $\beta_1(\text{PV}_{\text{supp}})$ | Effect in L5 of PV+ suppression | 0.0198 | [0.0068 0.0328] | 0.0028 |
| $\beta_2(\text{L2/3})$ | L2/3 vs. L5, with no light | 0.0181 | [0.0100 0.0262] | <0.0001 |
| $\beta_3(\text{SOM}_{\text{supp}}, \text{L2/3})$ | Interaction between SOM+ suppression and L2/3 recording | 0.0018 | [-0.0011 0.0047] | 0.22 |
| $\beta_3(\text{PV}_{\text{supp}}, \text{L2/3})$ | Interaction between PV+ suppression and L2/3 recording | 0.0246 | [0.0203 0.0290] | <0.0001 |
| DURATION | *Intercept* | 44.3 | [41.7 47.0] |  |
| $\beta_1(\text{SOM}_{\text{supp}})$ | Effect in L5 of SOM+ suppression | 1.10 | [1.02 1.28] | 0.012 |
| $\beta_1(\text{PV}_{\text{supp}})$ | Effect in L5 of PV+ suppression | 0.85 | [0.79 0.92] | <0.0001 |
| $\beta_2(\text{L2/3})$ | L2/3 vs. L5, with no light | 0.82 | [0.75 0.89] | <0.0001 |

Almost all spikes from single-cell recordings occurred during bursts, but occasionally some cells spiked before bursts. We presume that pre-burst spiking activity is necessary to initiate bursts, though we found that this activity was very sparse compared to participation in the bursts themselves, as we have shown previously (Krause et al. 2014). The probability of firing at least one spike before a burst (“early spikes”) was actually lowest in pyramidal cells and highest in PV+ cells (Figure 2F; 9/93 pyramidal, 6/28 SOM+, 12/29 PV+). Even among the special subset of cells that fired some early spikes, most spikes occurred after burst onset (Figure 2G). Very few early-spiking cells were in supragranular layers (Figure 2H); most early-spiking PV+ cells were in layer 4, whereas most early-spiking pyramidal and SOM+ cells were in layer 5 (Figure 2H). These results are consistent with previous reports of robust thalamic excitation of granular layer PV+ cells (Pouille and Scanziani 2001; Rose and Metherate 2005), the relative density of infragranular cell spiking (Barth and Poulet 2012; Krause et al. 2014; Sakata and Harris 2009), and direct thalamocortical excitation of infragranular cells (Constantinople and Bruno 2013; Krause et al. 2014; Tan et al. 2008).

### Timing of spikes relative to stimulus train

Sensory information in vivo is likely to be conveyed during high conductance states manifested as desynchronized or so-called UP states (Destexhe et al. 2007; Destexhe et al. 2003; Harris and Thiele 2011). During these periods of elevated network activity, high levels of synaptic input and net depolarization of cellular membrane potential will impact substantially on the precision and reliability of spike timing (Destexhe et al. 2003; Pachitariu et al. 2015), but there have been few systematic studies of spike timing and its control by GABAergic interneurons during such periods of high network activity. We used thalamically-induced network bursts as a model high conductance state, and assayed spike timing during these network bursts in response to ongoing thalamic stimulation.

The precision of spike timing relative to the TC stimulus train varied between and within cell types (Figure 3A-C). Some cells, especially PV+ interneurons, exhibited firing that was tightly linked to the ongoing stimulus train and relatively independent of the induced network activity (e.g. cell 9, Figure 3C). However, in most cells precision of spike timing relative to thalamic stimulation was poor (e.g. cells 1 & 2, Figure 3A). This finding was surprising. Our stimulus paradigm consisted of precise, synchronous afferent inputs. Spike timing relative to auditory stimulus features is very precise earlier in the auditory hierarchy (Bartlett and Wang 2007; Krishna and Semple 2000; Langner 1992; Wang et al. 2008), and specializations in auditory cortex for rapid processing of incoming input suggest that timing information is also important in auditory cortex (Kayser et al. 2010; Rose and Metherate 2005).

**Figure 3.**
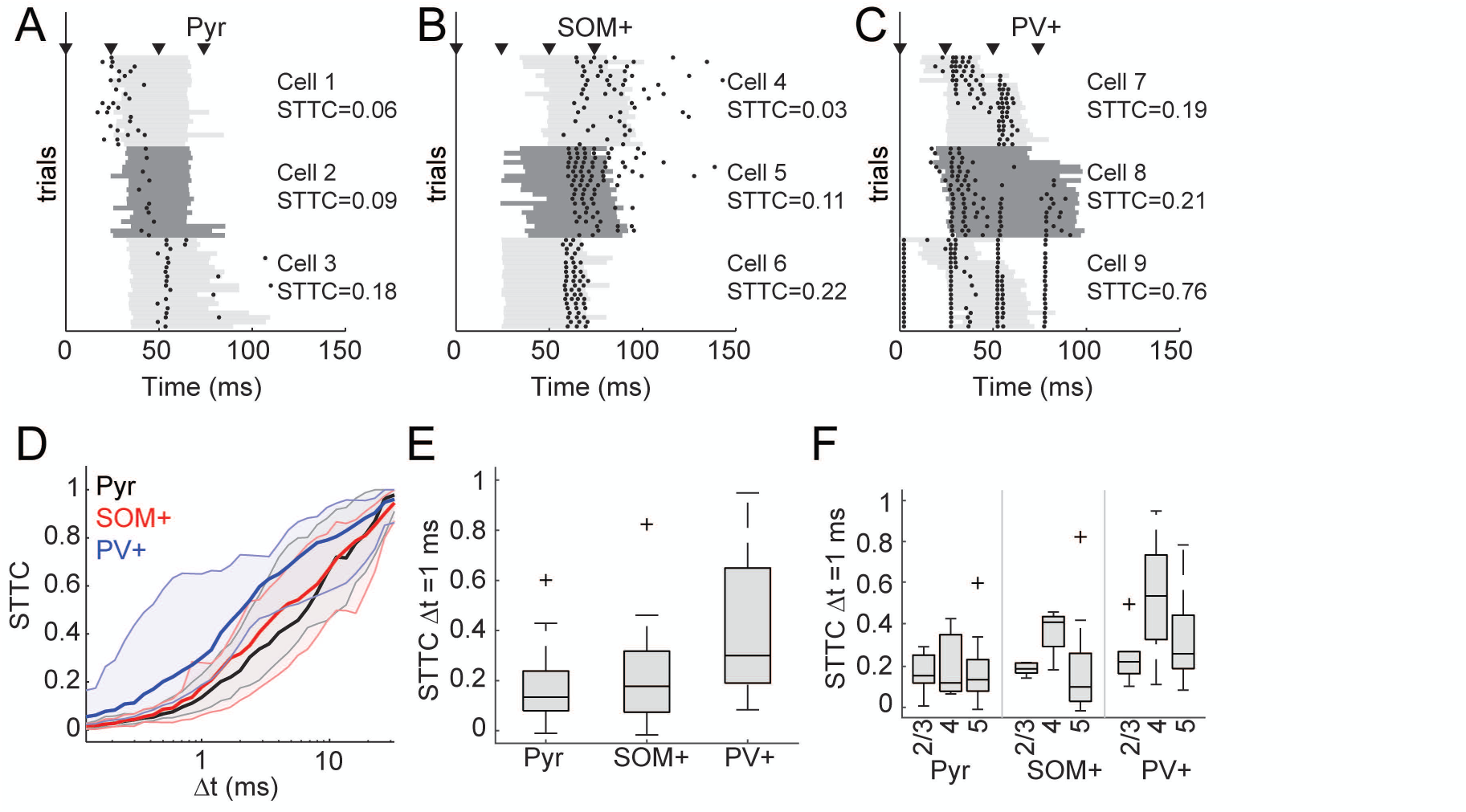
Spike timing with respect to stimulus. (A) Spike rasters for three example pyramidal cells, each with 20 trials. Bursts are indicated by gray bars that alternate shade for the three cells. The cells are plotted in order of increasing (from top to bottom) spike-time tiling coefficient (STTCs; see Methods). STTC values listed are for Δt = 1 ms. (B-C) Same as in (A), but for three SOM+ (B) and three PV+ (C) cells. (D) STTC plotted as a function of the time window (Δt) in which spikes were considered synchronous for pyramidal cells (black), SOM+ cells (red), and PV+ cells (blue). Dark lines are medians, and shaded regions indicate the interquartile range. (E) Comparison of STTC between cell populations at Δt = 1 ms. Spiking activity in PV+ cells was better timed across trials than for pyramidal or SOM+ cells. (F) The data in (E) separated by cortical layer. Overall, across all cell types cells in layer 4 were better timed (see Results).

To investigate further, we quantified stimulus-related spike timing in each cell class using the STTC (see Methods). A second measure, a variation on the Victor-Purpura spike train distance (Victor and Purpura 1996) (see Methods), gave nearly identical results (Supplementary Figure 4A). Importantly, we chose to use the STTC as a measure of spike timing because other measures like the cross correlation show increases with firing rate (Cohen and Kohn 2011; Cutts and Eglen 2014; De La Rocha et al. 2007). We calculated the STTC over a range of coincidence windows within which spikes were deemed coincident (Figure 3D). Although there was variation within cell types, PV+ cells tended to have a higher STTC over a wide range of coincidence windows. For precision of Dt = 1 ms (Figure 3E), PV+ cells were significantly better timed than both SOM+ cells (F(2,88)=12.73, p<0.0001; SOM+ vs. PV+ p=0.0014) and pyramidal cells (Pyr vs. PV+ p<0.0001). We also considered the effect of timing by layer (Figure 3F), which showed more precise firing in layer 4 for all types of cells, presumably reflective of larger TC EPSPs in this layer (Krause et al. 2014). A two-way ANOVA was significant for main effects of cell type (F(2,82)=6.42, p=0.0026) and layer (F(2,82)=4.26, p=0.017) but the interaction between cell type and layer was not significant (F(4,82)=0.86, p=0.49). The main effect of layer was driven by greater precision in layer 4 compared to the other layers (layer 2/3 vs. layer 4, p=0.023; layer 4 vs. layer 5, p=0.030).

**Figure 4.**
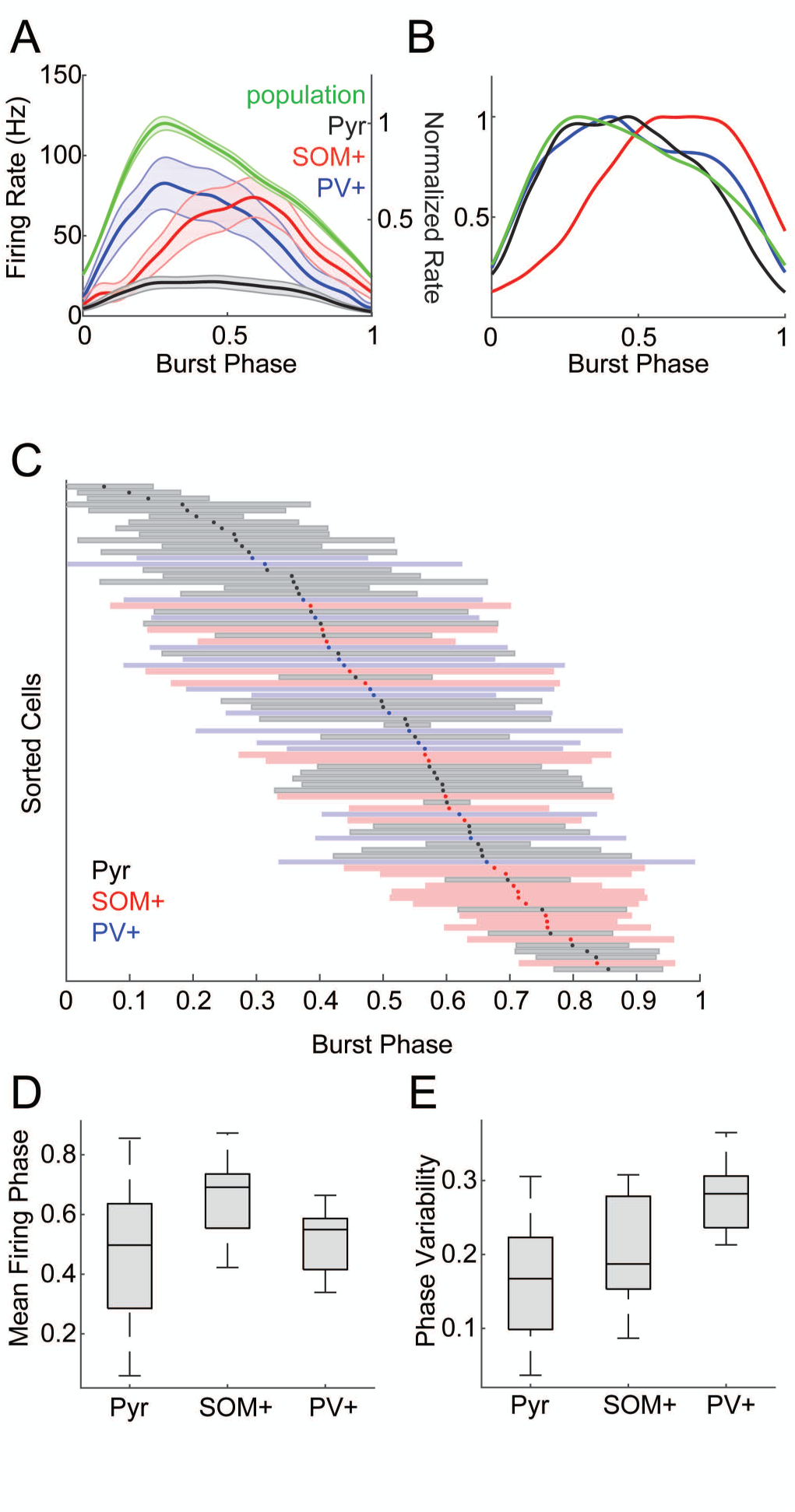
Single-cell effects of suppressing inhibition on firing rates and stimulus-related timing. (A) Spike probability for pyramidal cells recorded in slices with SOM+ or PV+ cells expressing inhibitory opsins in control conditions versus with inhibition suppressed. (B) As in (A), for spikes fired per trial. (C) Spike rasters for three example pyramidal cells, like in Figure 3A. (D) Suppressing SOM+ cells had little effect on these cells’ timing. (E) Same as (C), but for cells recorded in slices in which PV+ cells express inhibitory opsins. (F) Suppressing PV+ cells led to a pronounced improvement of pyramidal cells’ stimulus-related timing. (G) The spike time tiling coefficient, calculated at Δt=1ms, for cells recorded with and without optogenetic suppression of SOM+ or PV+ cells. (H) Effects on STTC were weakly correlated with effects on burst duration. In (A) and (G), * indicates significant differences based on a Wilcoxon signed-rank test (α=0.05).

### Stimulus-based timing is improved by suppressing PV+, but not SOM+ cells

The data presented so far indicate that PV+ cells are strongly and precisely driven by thalamic stimuli (Figure 3), suggesting that they are poised to provide precise feedforward inhibition in pyramidal cells. These observations are in line with previous reports, which have suggested that thalamically-evoked action potentials in cortical pyramidal cells are tightly regulated by inhibitory circuits (Cruikshank et al. 2007; Gabernet et al. 2005; Higley and Contreras 2006; Isaacson and Scanziani 2011; Oswald et al. 2006; Pouille and Scanziani 2001; Rose and Metherate 2005; Tiesinga et al. 2008; Wehr and Zador 2003). Specifically, these studies showed that inhibitory cells constrain integration windows and the timing of spikes in pyramidal cells, contributing to the information capacity of the cortical network. We sought to determine whether inhibitory cells similarly regulate spike timing in pyramidal cells during bursts. We expected that suppression of PV+ cells in particular would release pyramidal cells from inhibition, causing them to fire more indiscriminately and decrease the precision of spike timing relative to the stimulus. We tested this hypothesis by recording from individual pyramidal cells and measuring effects of optogenetic suppression of either PV+ or SOM+ cells on spike rate and timing. We suppressed SOM+ or PV+ cell activity by expressing halorhodopsin (via viral vectors) or ArchT (via transgenics) in SOM-Cre (n=43) and PV-Cre (n=23) mice (Supplementary Figure 3) and compared spike timing in pyramidal cells with and without suppression.

Suppression of either SOM+ or PV+ interneurons only modestly increased the firing rate of pyramidal cells during bursts. Pyramidal cell spike probability (Figure 4A) trended towards increasing for SOM+ cell suppression (Wilcoxon signed-rank test, z=1.86, p=0.062) and was significantly increased for PV+ cell suppression (z=3.66, p=0.0002). Pyramidal cells fired more spikes per trial (Figure 4B) with suppression of either cell type (SOM+ medians 0.76 versus 1.28 spikes/trial, z=2.46, p=0.014; PV+ medians 0.90 versus 1.06 spikes/trial, z=4.24, p<0.0001). Thus, pyramidal cell firing during bursts becomes slightly less sparse when SOM+ or PV+ cells are suppressed.

We expected that suppression of PV+ cells in particular would also lead to less precise spike timing in pyramidal cells relative to TC input. However, we observed no evidence of degraded timing of pyramidal cell firing during suppression of PV+ or SOM+ cells (Figure 4C-F). Contrary to our expectations, suppressing PV+ cells actually improved stimulus-related timing of most pyramidal cells (Figure 4G), an effect not observed with suppression of SOM+ cells. We observed the same result across a wide range of coincidence intervals, using the spike similarity measure, and in the cross correlogram (see Supplementary Figure 4B-D). There are several possible reasons for this unexpected result, though we were unable to identify one specific factor. Relief from strong feed-forward inhibition could have biased spiking in favor of responses to direct TC input rather than intrinsic network activity. Alterations in the dynamics of the bursts themselves may also play a role. Changes in STTC in pyramidal cells were weakly correlated with reductions in burst duration (Figure 4H; r^2^= 0.23, p=0.04)). We also observed on average a reduction in the standard deviation of burst onset latency, but this reduction only trended towards correlation with changes in STTC (r^2^= 0.16, p=0.10). These latter effects, which suggest a relationship between burst dynamics and timing, led us to investigate how PV+ and SOM+-mediated inhibition regulates spiking activity within network bursts. Below, we present data suggesting that PV+ cells do play a role in improving spike timing in pyramidal cells, but in the context of network activity rather than in the context of single cell responses to external inputs.

### Packet-like timing of spikes relative to network activity

Spike timing relative to an external stimulus can provide useful information, marking at a fixed latency the occurrence of stimulus features within a particular neuron’s receptive field. However, spike latencies can also vary with stimulus parameters, e.g. intensity, orientation or location (Panzeri et al. 2001; Shriki et al. 2012). Therefore, extraction of unambiguous information about a stimulus requires an internal reference signal, such as one provided by endogenous network activity (Kayser et al. 2009). Organization of spike times relative to other members of the local ensemble has been proposed to underlie population codes in the cortical network (Abeles et al. 1993; Luczak et al. 2013). We examined the organization of spiking within network bursts in slices by considering the phase at which spikes were fired, and examined how this organization depended on activity of GABAergic cells. Importantly, we made a distinction between the variation of burst firing phase within populations of cells (separated by cell type) versus the properties of individual cells within those populations.

Mean pyramidal and PV+ cell firing rates coincided with population activity levels throughout bursts, though pyramidal cells fired sparsely; firing in SOM+ cells was denser later in the burst (Figure 5A). These general trends are more evident in rate-normalized traces (Figure 5B). However, it is important to also consider the timing of individual cells within each population. Previous work has shown evidence for organization of network spiking activity into brief “packets,” reflecting a stereotyped sequence of activation of individual cells. Information within these packets can be carried both by the subset of cells participating in the packet and by the relative phase at which cells are active within the packet (Luczak et al. 2013; Luczak et al. 2015). Maximum information transfer requires maximum entropy in the repertoire of codes but minimum entropy in the code for a given stimulus. Therefore, an “ideal population” that utilizes timing within a packet to encode information has high variance in timing across the population but also high precision within each cell. That is, for a downstream neuron monitoring the output of this cortical network, optimally discriminable spike times of different potential inputs would have mean firing phases distributed across the burst (high phase variability across the population) but low phase variability for each individual cell. To distinguish between these critically different types of variance, we refer to the standard deviation of mean firing phases across the population as “phase diversity” and the standard deviation of firing phases within a cell as “phase variability.”

**Figure 5.**
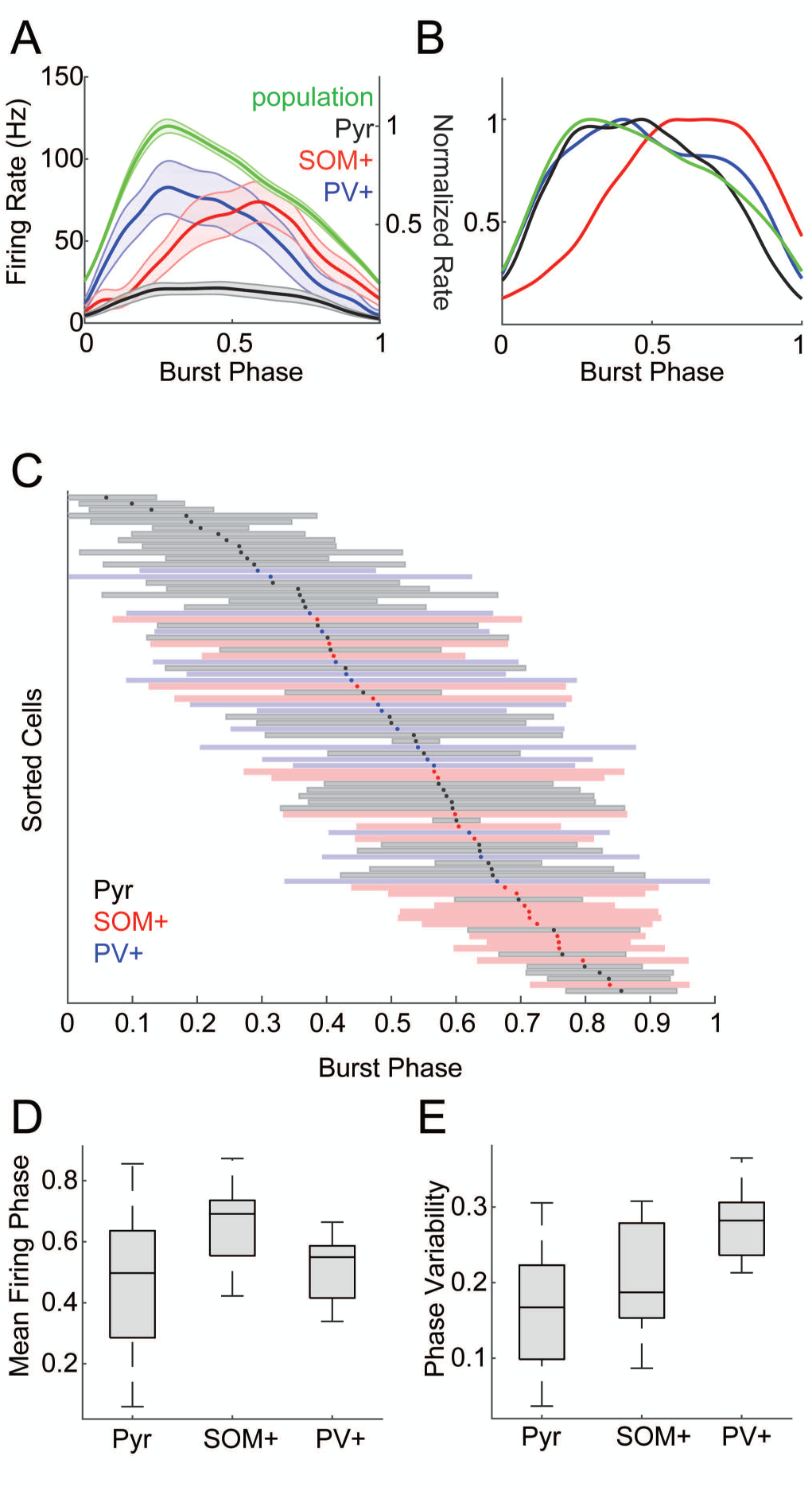
Spike timing with respect to bursts. (A) Firing rates averaged within populations as a function of burst duration for pyramidal (black), SOM+ (red), and PV+ (blue) cells. Population activity (green) is also plotted on an independent vertical scale, normalized to the peak population activity. Dark lines indicate means, shaded area is ±SEM. (B) Same as (A), but with all traces normalized to peak firing rate to illustrate the relative time courses of population activity in each cell type. (C) Timing of spikes across the duration of burst, normalized from onset (0) to offset (1). Cells are sorted by their mean firing phase relative to bursts, indicated with dots. Shaded bars indicate +/- one standard deviation. Color code indicates cell type: pyramidal cells, black/gray; SOM+ cells red, PV+ cells blue. (D) Mean firing phases for the data from (C). Phase diversity is indicated by the range of the distributions. Pyramidal cells had the most phase diversity. (E) Standard deviation of firing phase relative to bursts. Firing phase variability was lower in individual pyramidal cells compared to SOM+ or PV+ cells.

We computed the mean and variability of firing phase within the burst (defined as 0 at burst onset, 1 at burst offset) for each cell (Figure 5C). Over all cell types, firing phase spanned ∼80% of the burst duration, i.e. phase diversity was high, with a near uniform distribution of mean firing phases occurring during the central 60% of the burst. However, the distribution and diversity of mean firing phases and firing phase variability differed between cell types. Pyramidal cells had high phase diversity (Figure 5D), but the phase variability for individual pyramidal cells was low (Figure 5E), consistent with a population adapted for temporal coding. That is, that although pyramidal cell population activity was distributed throughout bursts, individual pyramidal cells tended to fire consistently at particular phases of the burst. By comparison, SOM+ and PV+ populations had less phase diversity (Figure 5D; Pyr vs. SOM+, Levene’s statistic (1,64)=8.51, p=0.0049; Pyr vs. PV+, Levene’s statistic (1,61)=9.97, p=0.0025) and individual cells had more phase variability (Figure 5E; F(2,81)=20.66, p<0.0001; Pyr vs. SOM+ p=0.040, Pyr vs PV+ p<0.0001). Consistent with the mean firing rates in Figure 5A, the mean firing phase of the SOM+ population was later than both pyramidal and PV+ cells (F(2,81)=8.01, p=0.0007; SOM+ vs. Pyr p=0.0004, SOM+ vs. PV+ p=0.0027). These results are consistent with the organization of pyramidal cell firing observed *in vivo* in which these cells tend to fire at consistent latencies relative to ‘packets’ (Luczak et al. 2013). However, although interneurons (particularly PV+ cells) were well-timed to input patterns (Figure 3), they were not well-timed to the output packet structure.

### Interneurons exert powerful control over network activity

To investigate the extent to which SOM+ and PV+ interneurons regulate network activity and the temporal organization of pyramidal cell spiking during bursts, we paired TC stimuli with optogenetic suppression of SOM+ or PV+ cells via halorhodopsin or ArchT; effects of the two opsins did not differ significantly and were pooled. We measured population activity in both layer 5 and layer 2/3 (Figure 6AB), because our previous observation that some bursts consisted primarily of activity in layer 5 without activity in layer 2/3 (Krause et al. 2014) suggested differing dynamics. We analyzed three measures extracted from the population activity: burst latency (Figure 6C), the peak of the MUA signal (Figure 6D), and burst duration (Figure 6E). We fitted a linear mixed-effects model to the population activity measures (see Methods). Model coefficient estimates for the individual parameters are in Table 2; we based our interpretations of significance on the means and standard errors of these parameter estimates. In Figure 6C-E, we plot means and standard error averaged across slices for each combination of factors.

In control conditions, bursts were earlier in layer 5 compared to layer 2/3 (center inset, Figure 6A&B; Table 2; Figure 6C), consistent with our previous results and with other results that showed activity starting in deep layers and propagating to superficial layers (Beltramo et al. 2013; Chauvette et al. 2010; Krause et al. 2014; Stroh et al. 2013; Wester and Contreras 2012). Layer 2/3 bursts were more intense than layer 5 bursts but did not last as long (Table 2; Figure 6D,E).

Unsurprisingly, suppression of either population of inhibitory cells increased burst amplitudes, but there were differential effects on burst latencies and durations, as well as differing magnitudes of effects on layer 5 compared to layer 2/3 bursts.

Suppressing either SOM+ or PV+ cells reduced burst onset latency in layer 5 (Figure 6C), but this effect was much greater for PV+ suppression (ratio of the latency with PV+ versus SOM+ cells suppressed = 0.64, 95% CI [0.58 0.71], p<0.0001). Both PV+ and SOM+ suppression increased the peak intensity of bursts (Figure 6D), but the effect was larger for PV+ compared to SOM+ suppression (PV - SOM=0.016 mV, 95% CI [0.003 0.029], p=0.017). By contrast, suppression of SOM+ versus PV+ interneurons had opposite effects on network burst duration. Bursts were shorter when PV+ cells were inactivated (Figure 6E), similar to the effect of suppressing inhibition pharmacologically (Sanchez-Vives et al. 2010). By contrast, bursts were longer when SOM+ cells were inactivated (Figure 6E), suggesting that suppression of SOM+ cells produces more complex effects than simple reduction in inhibitory tone.

The most dramatic effects were observed in layer 2/3 following suppression of PV+ cells. There was a slightly greater reduction in latencies following SOM+ and PV+ suppression in layer 2/3 compared to layer 5. Because bursts likely initiate in layer 5, this result suggests that suppression of inhibition led to more rapid spread of bursts to layers 2/3. This effect is likely driven by either the greater intensity of bursts in layer 5 (demonstrated by increased peak MUA) hastening the spread of activation to the supragranular layers, or the reduced inhibition in the supragranular layers increasing responsiveness, or a combination of both factors. PV+ suppression also had a bigger effect on burst peak in layer 2/3 compared to layer 5 (Table 2). The interaction of PV+-suppression with layer was significantly greater than the interaction of SOM+-suppression with layer (PV – SOM=0.0228 mV, 95% CI [0.0177 0.0280], p<0.0001), and the interaction of SOM+ suppression with layer was not significant (Table 2). These results suggest that PV+ cells exert a dominant influence on activity in the supragranular layers compared to their influence in the infragranular layers and compared to the influence of SOM+ cells, consistent with the concentration of PV+ cells in upper layers of cortex (Supplementary Figure 2D), though both interneuron populations can influence the rate at which activity spreads between laminae.

### Suppressing PV+ cells impairs packet-based timing

Spike timing during network bursts can greatly enhance the amount of information carried in a population (Borst and Theunissen 1999; Contreras et al. 2013; Kayser et al. 2012; Kwag et al. 2011; Luczak et al. 2015; Panzeri et al. 2001). If information is encoded in temporal patterns of spikes, then the relevant timing information may not solely be timing relative to the stimulus (Figure 4), but also timing relative to population activity (Figure 5). We showed above that there is low variability in firing phase in individual pyramidal cells within bursts, but there is high diversity in firing phase across the pyramidal cell population (Figure 5), indicating pyramidal cells can encode information by their phase of firing (Kayser et al. 2009). Next, we show that interneurons play a role in organizing this network-based spike timing by repeating the analysis of Figure 5C with and without suppression of SOM+ and PV+ cells. Consistent with their more modest effects on burst properties illustrated in Figure 6, suppressing SOM+ cells had little impact on the order that pyramidal cells fired during bursts (Figure 7A-B) or on the mean firing phase during bursts (Figure 7E; mean phase change=0.007). Thus, as for spike timing relative to afferent stimuli (Figure 4G), SOM+ cells appear to exert little regulatory control over the temporal organization of pyramidal cell spiking during bursts. By contrast, suppression of PV+ cells altered the temporal sequence of activation (Figure 7C-D), primarily by causing some cells that fired later in bursts to fire earlier (Figure 7E; mean phase change=0.10, PV+ versus SOM+ effect t(33)=2.52, p=0.017), and phase diversity was significantly lower with PV+ suppression (Figure 7E; Levene’s statistic (1,38)=4.63, p=0.038). This loss of phase diversity compressed the fractional span of the burst over which the population of pyramidal cells was active, thereby reducing the capacity of a potential temporal population code. Suppression of neither PV+ nor SOM+ cells had a significant effect on consistency of burst firing phase in individual cells (Figure 7F).

**Figure 7.**
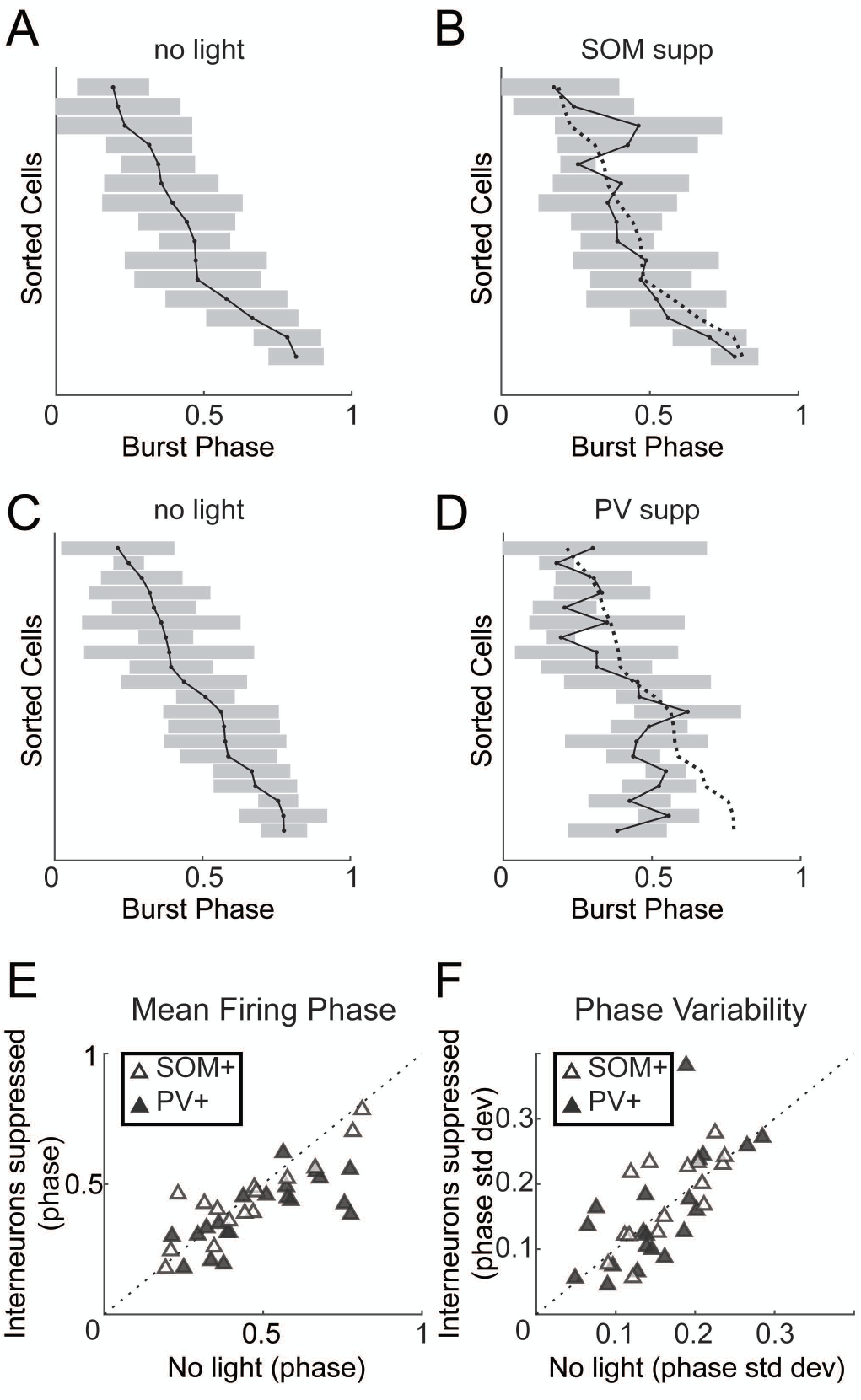
Effects of suppressing inhibition on timing with respect to bursts. (A) Burst firing phase means (points) and ±1 standard deviation (shaded) for pyramidal cells. (B) Same cells as in (A) but with SOM+ cells suppressed. Dotted line in (B) shows the original firing phase means from (A). (C-D) Same as (A-B), but showing how suppression of PV+ cells (D) altered the temporal sequence of firing. (E) Mean firing phase of pyramidal cells relative to bursts. Suppression of PV+ cells caused pyramidal cells to fire earlier in bursts and reduced phase diversity. (F) Phase variability is not significantly affected by suppressing either interneuron population.

We observed two effects of suppressing PV+ cells that could potentially impact the organization of firing patterns during network bursts. First, the bursts themselves became shorter (Figure 6E), compressing the temporal scale of spike packets during bursts. Second, the diversity of spike phases available to the network during a burst was reduced with suppression of PV+ cells, compressing the available time scale for population-based codes (Figure 7D). Suppression of PV+ cells also (modestly) increased firing rates of pyramidal cells (Figure 4A,B) and had differential effects on network activity in supragranular versus infragranular layers (Figure 6). Because it was unclear how each of these changes might impact the ability of the network to produce distinct patterns of activity critical for population codes, we implemented a simple decoding model that interprets spike patterns, the tempotron (Gütig and Sompolinsky 2006). The tempotron is an integrate-and-fire model that produces a binary decision (spike or no spike) in response to patterns of synaptic inputs. Our main goal was to test the ability of the model to distinguish temporal patterns within network activity based on the structure of such activity observed in control conditions versus with PV+ cells suppressed. The model was driven by simulated spiking activity of 100 units with firing statistics based on our recordings from pyramidal cells during bursts (Figure 8A, top). The tempotron integrates these inputs over time and produces a “spike” on trials in which a threshold is reached (Figure 8A, right).

**Figure 8.**
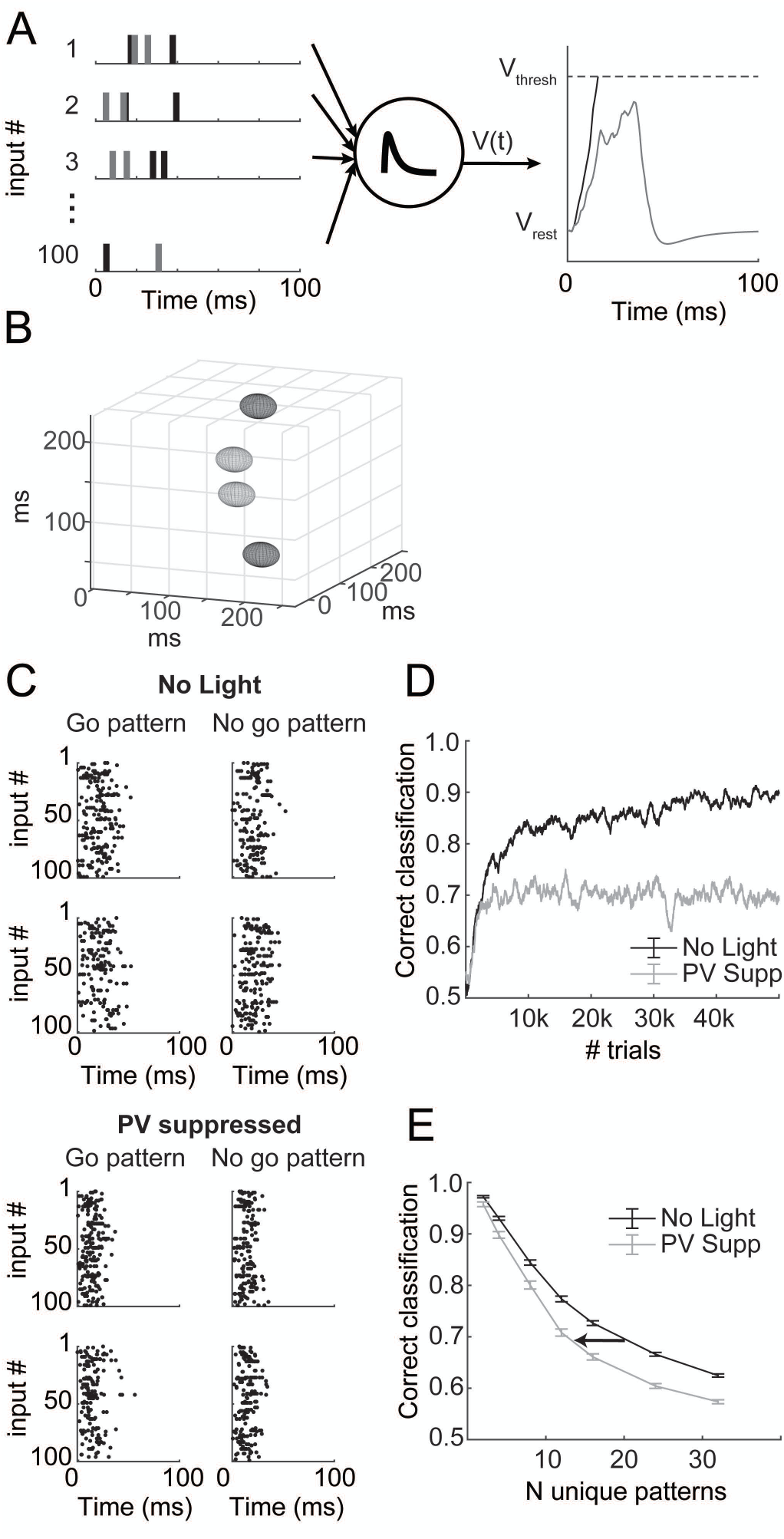
Decoding simulated patterns in bursts. (A) Input from 100 spike trains (left) is summed by a leaky integrate-and-fire unit (center) called a tempotron. On a given trial, this unit produces a spike if it reaches threshold (right, black), otherwise it produces no spike (grey). (B) Input patterns depicted as 3D ellipsoids, representing the mean (position) and phase variability (semi-axes of ellipsoids) for three of the 100 total input units. The semi-axes represent only 1/10 of one standard deviation; the patterns are quite overlapped in three dimensions compared to the simplified toy representation here, which is why many inputs are necessary and why performance is not perfect. The “go” (light gray) and “no go” (dark gray) patterns do not have any spatial organization in this space. (C) Randomly generated input spike rasters for two example “go” and two “no go” patterns, for simulations based on “no light” experiments (top) and based on PV+-suppression experiments (bottom). Note that there are no obvious similarities among the go versus no go patterns, and that the patterns based on PV+-suppression data are compressed in time. (D) As the tempotron learns by adjusting the weights of the input units (see Methods), performance increases but eventually plateaus. These traces represent a single learning example. (E) Mean performance (±standard error) after 50,000 trials for 100 unique model runs at each point indicated. The number of unique patterns indicates the difficulty of the classification. Simulations based on PV+-suppression underperform compared to control results at all difficulty levels.

In these simulations, we generated a set of 2 – 32 stochastic spike patterns with statistics that mimic recorded pyramidal cell activity. Patterns differed in that the mean firing phase for each input unit was randomized between patterns. The mean firing phase for each unit and pattern was selected from a smoothed distribution based on observed data (Figure 5D). The firing rate and variance were for each unit were also randomly selected from a smoothed distribution based on recorded data (Figure 5A; Figure 7F), but these parameters were kept constant across patterns. Burst durations were the mean burst durations from experiments in Figure 6. Random training and testing trials were generated using these simple statistics for each input unit. Because of these parameter choices, patterns can be thought of as arbitrary clouds of spike patterns in an N-dimensional space, where N is the number of input units; a toy example with 3 units is illustrated in Figure 8B. The size in each dimension reflects the variance in firing phase for each unit, creating an ellipsoid. Because the variances are constant across patterns, but the mean firing phases are not, each pattern is represented by an ellipsoid of constant shape but varying position (four patterns are illustrated in the example of Figure 8B). “Correct” classification was deemed as “go” (i.e. the tempotron should fire a spike) for half of the patterns and “no go” (i.e. no spike) for the others (see Supplementary Figure 5). Classifying these patterns is difficult, because there is no predetermined structure of those go/no go patterns in the N-dimensional space nor any non-random similarity among the “go” patterns or among the “no go” patterns, and because the variance of each contributing input unit is large. (In Figure 8B, we depict each ellipsoid with semi-axes representing 1/10 of 1 standard deviation so the patterns can be easily distinguished by eye.)

In Figure 8C (top) we show spike rasters for four example patterns across all 100 input units under control conditions (top four panels) and with PV+ suppression (bottom four panels). The biggest effects of PV+ suppression on patterns were that bursts were shorter and that activity was earlier within the burst, resulting in temporal compression of patterns in PV+-suppressed conditions (Figure 8C, bottom).

The tempotron was trained by altering synaptic weights (see Methods) according to responses to the patterned input (Figure 8D). As the number of input patterns increases, the classification task becomes more complex, leading to poorer steady-state performance (Figure 8E). Training within the control-based and PV suppression-based populations was completely independent.

In these simulations, we tested the discriminability of patterns generated by populations of units with different relative temporal and rate statistics observed under control conditions and with PV+ cells suppressed. We assayed the ability of the control versus PV suppression-based populations to encode temporal information by the performance of the tempotron after 50,000 training trials for a range of pattern counts (i.e. discrimination complexities). For a given pattern count, inputs from control based populations modestly outperformed those from PV-suppressed populations (Figure 8E). However, we also found that there was a large impact on the potential complexity that could be represented at a moderate performance level (for example, 70%) (Figure 8E; arrow depicts a leftward shift towards less complexity). Therefore, we conclude that PV-mediated inhibition may allow networks of pyramidal cells to encode a greater number of distinct patterns by increasing the duration of network bursts.

A further advantage of this modeling approach is that we can assay the effects of individual parameter changes by choosing which effects to incorporate into the model. The two effects of PV-suppression most likely to contribute to impaired discrimination performance are the reduced duration of bursts (Figure 6E) and the reduced phase diversity across the pyramidal cell population (Figure 7D,E). These effects jointly contribute to the temporal compression seen in Figure 8C. To determine whether one of these factors was the dominant influence on impaired discrimination in the PV-suppressed population, we performed simulations using PV-suppressed population statistics except we substituted either control burst durations (while maintaining PV-suppressed firing phase statistics) or control mean firing phase distributions (while maintaining PV-suppressed burst durations) into the model. Either manipulation partially relieved the temporal compression caused by PV-suppression. We found that using control burst durations only recovered the PV-suppression effect slightly, but using control mean firing phase distributions completely recovered the PV-suppression effect (Figure 9A). Therefore, we conclude that while both of these factors affect encoding potential, the reduced range of firing phases, i.e. the concentration of mean firing phase early in bursts, was a key mediator of the effect of suppressing PV+ cells.

**Figure 9.**
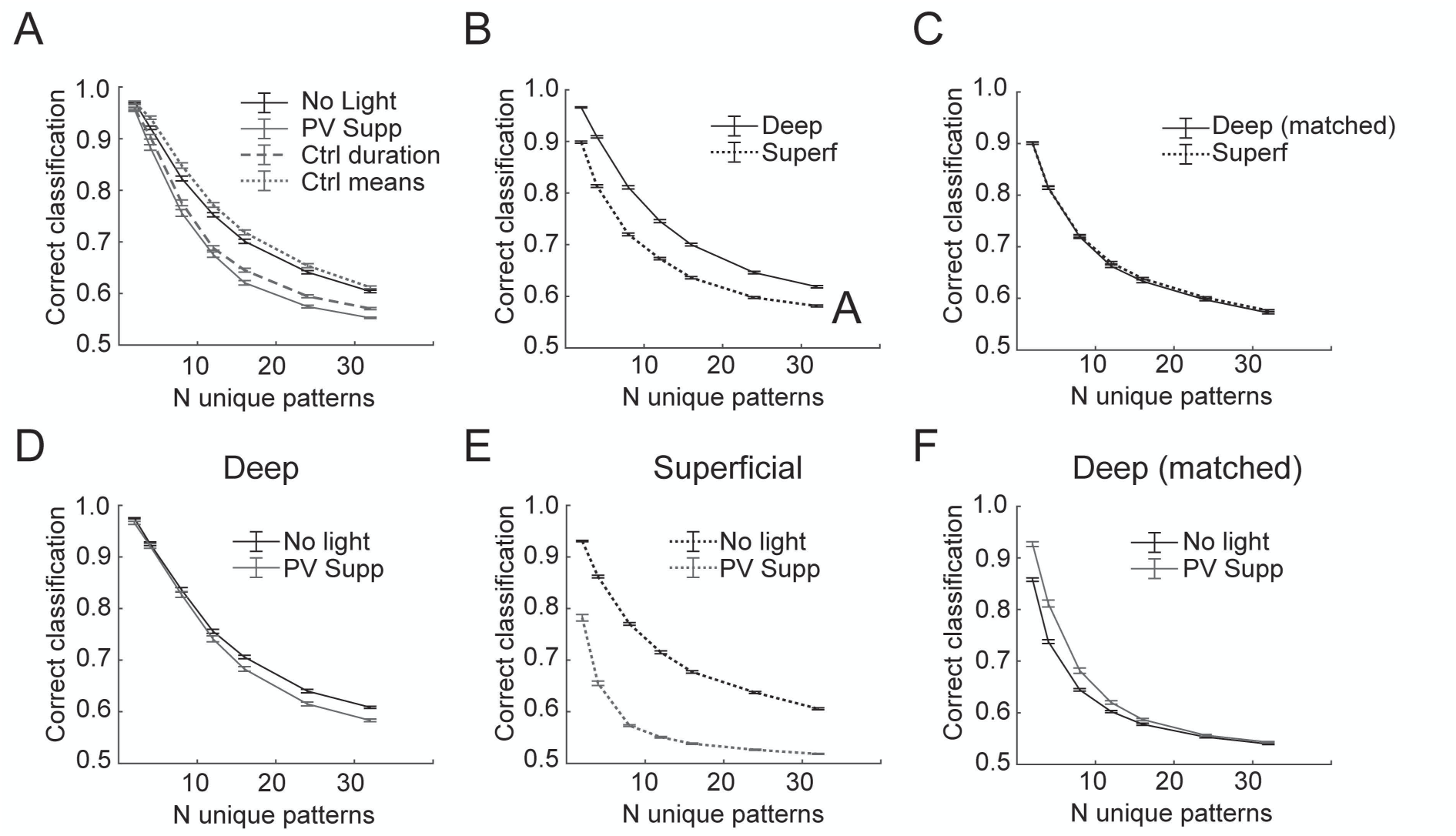
Parameter and laminar influences on decoding. (A) Results from Figure 8E (solid lines) compared to models that used mostly PV+ suppression parameters except for burst duration (dashed line) or mean distribution (dotted line). (B) Model performance using deep (solid line) versus superficial (dotted line) pyramidal cells as inputs from no light condition. (C) Model performance using firing rates for deep cells matched to the firing rates of superficial cells. (D-E) Effects of PV+ suppression on model performance for models using only deep (D) or only superficial (E) inputs. (F) Effects of PV+ suppression on performance with deep cell inputs matched to the firing rates of superficial cells. Error bars represent ±SEM for all panels.

Although superficial and deep-layer cells can act in one functional circuit in a cortical column, pyramidal cells are most highly connected with other cells in the same layer, and populations of cells from different layers have different long-range projection patterns. Differences in tuning properties and firing density in superficial and deep pyramidal cells likely contribute to differences in sensory coding (Barth and Poulet 2012). Our simulation results emphasize the importance of firing rate during bursts for population coding capacity. Thus, the tempotron performed better with inputs whose firing statistics were based on recordings from deep cells under control conditions than with inputs based on superficial cells (Figure 9B). However, there was no longer a difference between populations if we subsampled the deep cells to produce mean firing rates comparable to superficial cells (Figure 9C).

We also show in Figure 6 that PV- suppression appears to have the most profound effects in the superficial layers. Therefore, we also used the tempotron model to separate the effects of PV-suppression on deep versus superficial cells. Our simulation results suggest that the effects on population coding of suppressing PV+ cells were manifested primarily in supragranular layers. We found that PV-suppression actually had very little effect on tempotron performance using the deep population (Figure 9D), but had a substantial effect using the superficial population (Figure 9E). The differential effect was not due to differences in firing rate, because the deep population was still insensitive to PV-suppression when we resampled to match to superficial firing rates (Figure 9F). These simulations suggest that PV+ cell-mediated inhibition in supragranular layers strongly regulates the information encoding capacity of the local cortical network, consistent with its greater effects on network activity in supragranular layers and with reports suggesting that sparse firing in supragranular cells contributes to their importance in sensory information processing (Barth and Poulet 2012).

## Discussion

### Summary

We used burst activity induced by TC afferents to auditory cortex as a model to study the spatial and temporal structure of correlated network activity in the cortical column and its regulation by local GABAergic interneurons. We note that in this preparation, afferent stimuli served both to initiate network activity and as a model for ascending sensory information conveyed to cortex on the background of ongoing activity during the desynchronized state. Three distinct cell classes, pyramidal cells, PV+ cells and SOM+ cells, exhibited differences in density and timing of spiking activity during these network bursts and relative to patterned afferent stimulation. The precisely timed spikes observed in PV+ cells in response to TC stimuli suggest they are poised to regulate spike timing in pyramidal cells via feedforward inhibition, as observed in somatosensory cortex (Gabernet et al. 2005). In contrast to these expectations, however, spike timing in pyramidal cells was poor under control conditions and improved upon optogenetic suppression of PV+ cells. Instead, our data suggest that the organization of spiking during network bursts is consistent with observations *in vivo* of population codes relying on packet-based stimulus representation. Furthermore, our data suggest that PV+ cells act to regulate spike timing within these bursts by maintaining this temporal structure. Simulation results suggest that the temporal sequence of activity within the pyramidal cell population is important for producing population spike patterns that can be categorized by downstream neurons, and suppressing PV+ cells degrades the ability of the network to produce discriminable activity patterns. This effect is most pronounced for supragranular pyramidal cells, consistent with previous results indicating a critical role for sparsely-firing supragranular cells in sensory coding (Barth and Poulet 2012; Crochet et al. 2011; DeWeese et al. 2003; Hromadka et al. 2008; Poulet and Petersen 2008; Sakata and Harris 2009).

### Spike timing in auditory cortex

The ability of pyramidal cells to encode stimulus information depends not only on firing rates but also on precision and reliability of responses across trials (Kayser et al. 2010; Mainen and Sejnowski 1995). Because PV+ cells have been shown to constrain pyramidal cell spikes to particular windows of time (Cruikshank et al. 2007; Gabernet et al. 2005; Pouille and Scanziani 2001; Rose and Metherate 2005; Wehr and Zador 2003; Zhu et al. 2015), we expected that suppressing PV+ cells would impair pyramidal cell timing. Furthermore, spiking responses to TC afferents in PV+ cells were better timed than in other cell types (Figure 3), positioning them to precisely control spike timing.

These expectations were unmet in our observations in two ways. First, pyramidal cell spiking responses to afferent stimulation were not well timed (Figure 3). Previous work (Gil et al. 1999; Krause et al. 2014; Rose and Metherate 2005) has shown that TC EPSPs are precisely timed and large, especially in layer 4 pyramidal cells, suggesting that spikes would be precise and readily evoked. The absence of precise timing in pyramidal cells relative to the stimulus train was unexpected given the importance of, and specialization for, timing information in the ascending auditory pathway (Elhilali et al. 2004; Heil and Irvine 1997; Phillips and Hall 1990; Rose and Metherate 2005), but may reflect a transition from timing/feature-based coding to rate/object-based coding in auditory cortex (Wang et al. 2008). Indeed, we have shown that monosynaptically driven spikes in pyramidal cells are rare in auditory cortex, with most spiking occurring in the context of network bursts induced by thalamic stimulation (Hentschke et al. 2017; Krause et al. 2014). On the background of this ongoing activity, thalamic synaptic responses are less effective at driving precise spiking responses, and spike timing may be influenced significantly by intrinsic cortical activity. This result is consistent with degraded spike timing information in auditory cortex compared to the periphery (Chechik et al. 2006; Linden et al. 2003; Nelken 2004; Ter Mikaelian et al. 2007), even though first-spike precision persists in many cells (Heil and Irvine 1997; Phillips and Hall 1990).

Second, suppression of PV+ cells did not further degrade spike timing in pyramidal cells. In fact, spike timing was improved in the absence of strong PV+ cell-mediated inhibition (Figure 4). The basis for this improvement is not entirely clear. It is possible that PV+ cell-mediated feedforward inhibition was initially strong enough and fast enough to prevent direct (monosynaptic) TC input from producing spikes in most cells. That is, if we consider input to pyramidal cells as arising from either the direct (monosynaptic) TC input or from cortico-cortical inputs associated with the network burst, suppressing PV+ cells could bias the pyramidal cells to respond to direct TC input due to disruptions in the balance of excitation and inhibition (Gabernet et al. 2005; Isaacson and Scanziani 2011; Vogels and Abbott 2009; Wehr and Zador 2003). Alternatively, it is possible that changes in the properties of bursts themselves underlie this effect. For example, there is a negative correlation between the reduction in burst duration and the improvement in STTC and a potential correlation with decreased variability (Figure 4). This suggests that briefer, more stereotyped bursts cause an apparent increase in spiking precision. Whatever the basis for the improvement, it is clear that we did not observe the expected degradation in spike timing upon suppression of PV+ cells.

### Network activity in auditory cortex

Classical studies of response properties in sensory cortex characterized single cells in terms of tuning curves, reflecting mean firing activity across many trials (Hubel and Wiesel 1959; Spillmann 2014). Much of what we understand about topography and hierarchical processing in cortex is based on these representations. Implicit in these analyses, however, is the assumption of independent firing in individual cells. Population codes in these models are derived from assaying spike rate or timing based on each cell’s tuning curve for the stimulus of interest. However, the importance of correlated spiking activity in cortical networks that arises due to intrinsic connectivity and shared afferent input has long been recognized as well (Kohn et al. 2009; Panzeri et al. 1999). Data suggest that local cortical networks, and in particular the cortical column, can operate as a unit with all-or-none responses to afferent stimulation analogous to the behavior of single neurons (Bathellier et al. 2012; Luczak et al. 2015; Yuste 2015).

Correlated spiking activity in neocortex arises in a variety of circumstances, but is particularly prominent when the TC network is in the ‘synchronized state’. Here, the network cycles between ON and OFF periods of spiking activity and quiescence, respectively, and individual cells cycle between depolarized/high conductance and hyperpolarized/low conductance resting states (UP and DOWN states). The synchronized state is most obviously observed in slow wave sleep and under anesthesia, but can also be observed during quiet wakefulness, when the network can exhibit UP/DOWN transitions or dwell in extended DOWN periods (McGinley et al. 2015; Mochol et al. 2015; Petersen et al. 2003; Poulet and Petersen 2008). Upon sensory arousal and motor activation, the network eschews synchrony in favor of an extended ON period (desynchronized state), and, intracellularly, extended UP states. In auditory cortex, the transition from quiet wakefulness to arousal and especially locomotion has similar effects on network activity, with cells moving from ON/OFF cycling to extended ON periods (Zhou et al. 2014). However, evidence suggests that optimal stimulus detection and discrimination occurs during moderate arousal (McGinley et al. 2015). Here, activity is low and bursts are inducable, in contrast to periods of high arousal, when the network is depolarized and desynchronized, or low arousal, when the network exhibits spontaneous UP-DOWN state transitions. This is consistent with the intuition that maximal sensitivity in audition occurs during quiet, alert periods.

Sensory stimulation triggers synaptic and spiking activity that occurs on the background of spontaneous network activity. The largest responses occur when activation of thalamic or cortical afferents triggers correlated network activity within the cortical column (Hentschke et al. 2017; Krause et al. 2014; Luczak et al. 2013; Sakata and Harris 2009). The spatio-temporal patterns of spiking activity (packets) during these network bursts in auditory cortex are preserved across the synchronized-desynchronized state continuum, suggesting that the fundamental structure of network activity can be elucidated by examining network bursts in isolation. Stimulus-related variability in spike packets during networks bursts is postulated to underlie population codes for sensory information (Luczak et al. 2015), while other neurons may provide a reliable timing signal to mark actual temporal structure (Brasselet et al. 2012).

The structure of network events we observe in auditory TC slices is similar in many regards to the structure of activity observed during sensory processing in auditory cortex *in vivo*. The firing rates of single cells during bursts (Figure 2) were comparable to those observed in auditory cortex *in vivo* (Hromadka et al. 2008; Luczak et al. 2013). The duration of network bursts we observe (Figure 1), is similar to the ∼50-100 ms “packets” (Luczak et al. 2013) or ∼50 ms “bumps” (DeWeese and Zador 2006) observed in auditory cortex *in vivo*. Unlike in auditory cortex, spontaneous and evoked bursts or “UP states” in other cortical areas tend to last for hundreds of milliseconds to seconds (Neske et al. 2015; Sanchez-Vives and McCormick 2000). These data suggest that the short duration of bursts in auditory cortical columns reflect the operation of auditory cortical networks on rapid timescales, consistent with other temporal specializations in the ascending auditory pathway (Elhilali et al. 2004; Heil and Irvine 1997; Phillips and Hall 1990). Furthermore, the spatiotemporal structure of network bursts is comparable as well. Both *in vivo* and in slices burst activity originates in layer 5 and spreads to other layers (Beltramo et al. 2013; Chauvette et al. 2010; Krause et al. 2014; Sakata and Harris 2009; Wester and Contreras 2012). Variable involvement of supragranular cells, presumably due to their more hyperpolarized resting potentials during DOWN states as well as strong PV+-mediated inhibition, is observed both in slices (Krause et al. 2014) and *in vivo* (Sakata and Harris 2009). The temporal structure of pyramidal cell spiking activity we observe in slices, including mean firing phases that span almost the entire burst duration and consistent firing phase in single cells (Figure 5), is compatible with the packet-based organization of spiking activity observed *in vivo* (Luczak et al. 2015).

Overall, these similarities suggest that the experimental model used here to elucidate the role of inhibitory cells in regulating network activity and spike timing in pyramidal cells has direct implications for understanding cortical sensory processing *in vivo*. However, we note that reduced preparations exhibit substantially different activity patterns from those observed *in vivo*, likely reflecting the absence of subcortical neuromodulators and reduced numbers of synaptic inputs in the cortical column. Long range connections especially are lost in slice preparations, resulting in a significantly altered balance between excitation and inhibition (Stepanyants et al. 2009). These changes in connectivity result in reduced levels of spontaneous activity, and especially of spontaneous network bursting, which may preclude observation of richly varied network-level interactions observed *in vivo* (Kato et al. 2017; Seybold et al. 2015). Because of reduced excitatory inputs in slices, manipulations of inhibitory cells may overemphasize their overall contribution to network-level phenomena. On the other hand, because axonal projections of SOM+ cells tend to be more extended compared to PV+ cells (Caputi et al. 2013; Markram et al. 2004), it is possible that the experiments presented here have underemphasized their importance. Subsequent studies investigating spike timing during network activity *in vivo* will provide a more complete picture of how inhibitory and excitatory cell populations interact to produce firing patterns critical to sensory coding,

### Regulation of network activity by GABAergic cells

Both PV+ and SOM+ cells are strongly activated directly by afferent inputs and during network activity triggered by these inputs (Figure 2). Suppressing either cell type reduced burst onset latency and increased the peak intensity of bursts (Figure 6), but both of these effects were much greater for PV+ cells than SOM+ cells. These data suggest that PV+ cells exert far greater control over network activity compared to SOM+ cells, consistent with previous studies of regulation by UP state activity by GABAergic cells (Fanselow and Connors 2010; Neske et al. 2015). The opposite effect on duration of suppression of SOM+ (increased duration) versus PV+ cells (decreased duration) likely in part reflects differences in inhibitory tone during network bursts. That is, eliminating a major source of peri-burst inhibition by suppressing PV+ cells leads the network to burst intensely but dissipate at a faster rate (e.g. due to depletion of vesicles), while suppressing a less dominant form of inhibition in SOM+ cells allows burst activity to proceed for a longer period of time at only slightly elevated intensity. These effects are also consistent with the distinct temporal profiles of spiking activity of these cells during bursts (Figure 5). PV+ cells tended to be active before and throughout bursts, while SOM+ cells were active late. The combination of reduced latency and duration observed with suppression of PV+ cell is reminiscent of effects observed with moderate concentrations of GABAergic antagonists. These studies also showed an inverse relationship between burst intensity and duration (Sanchez-Vives et al. 2010), indicating that the relationships found with GABAergic antagonists reflect the influence of PV+-mediated inhibition.

The observations that the vast majority of spiking occurs in the context of network bursts (Krause et al. 2014), and that sensory stimuli induce burst-like responses from hyperpolarized states (Curto et al. 2009; McGinley et al. 2015; Sakata and Harris 2009), indicate that the overall gain of the cortical column is reflected in the magnitude of network bursts (Womelsdorf et al. 2014), which is controlled by activity of GABAergic cells (Figure 6). *In vivo*, modulation of PV+ cell activity modulates the gain of sensory responses (Atallah et al. 2012; Wilson et al. 2012) consistent with our observations that suppressing PV+ cells increases the magnitude of network bursts. Interestingly, although active behavior and attention are associated with increases in gain (Andersen and Mountcastle 1983; McAdams and Maunsell 1999; Salinas and Thier 2000; Treue and Trujillo 1999), attention can also increase firing rates of fast-spiking cells specifically, especially when tasks are difficult (Chen et al. 2008; Mitchell et al. 2007), suggesting instead a reduction in gain. These results and our observations are consistent with a model in which there is an optimal level of PV+-mediated inhibition depending on the type of task, where PV+ cell-mediated inhibition produces longer bursts of activity that allow for downstream circuitry to extract more nuance necessary for sensory discrimination on difficult tasks.

In contrast to our observations (Figure 5), SOM+ cells are far less active during bursts in somatosensory and entorhinal cortices compared to PV+ cells and have been proposed to play less significant functional roles in regulating network activity in those areas (Neske et al. 2015; Tahvildari et al. 2012). However, suppression of SOM+ cells substantially increased pyramidal cell firing rates in somatosensory cortex (Neske and Connors 2016), again in contrast to our results (Figure 4). Although we saw higher firing rates in SOM+ cells in auditory cortex, we still observed that SOM+ cells fired less precisely and reliably from trial to trial compared to PV+ cells, despite large TC EPSPs (Table 1). This result may be due to slower membrane time constants, lack of inhibition from PV+ cells to constrain spike times, or variable burst firing patterns triggered by afferent stimulation (Kawaguchi and Kubota 1996). Unlike previous studies (Fanselow and Connors 2010; Neske et al. 2015), we found that SOM+ cell activity is biased towards the end of bursts; we note that because these bursts are so brief in auditory cortex, “late” activity in our bursts may correspond with early activity in those other studies. In addition, SOM+ cell activity may depend on age (P28-56 here versus P12-18 in (Fanselow and Connors 2010; Neske et al. 2015; Tahvildari et al. 2012)) or how bursts are triggered (spontaneous or induced with cortical stimulation versus bursts induced by trains of TC input).

Due to facilitating excitatory inputs from pyramidal cells, SOM+ cells are well-positioned to contribute to burst termination (Krishnamurthy et al. 2012; Melamed et al. 2008; Reyes et al. 1998; Silberberg and Markram 2007). However, most evidence suggests that activity-dependent potassium conductances are more important than inhibition in terminating UP states (Compte et al. 2003; Hill and Tononi 2005; Neske 2016; Sanchez-Vives and McCormick 2000). We show here that suppression of SOM+ cells increases burst duration (Figure 6). Although the increase in duration was modest, the intensity and duration of bursts is typically inversely correlated, as seen in the optogenetic suppression of PV+ cells here and with GABAergic antagonists previously (Sanchez-Vives et al. 2010). Thus, the cooccurrence of increased duration and increased intensity observed with suppression of SOM+ cells is even more striking. The high level of SOM+ cell activity during bursts in auditory cortex may be a specialization contributing to the brevity of network bursts in auditory cortex, possibly via activation of GABA_B_ receptors. GABA_B_, but not GABA_A_, antagonists, can prolong UP states (Mann et al. 2009), and SOM+ cells trigger GABA_B_-mediated suppression of glutamatergic synapses (Urban-Ciecko et al. 2015).

The robust activity of SOM+ cells during bursts in auditory cortex may suppress feedback inputs to distal dendrites of pyramidal cells during sensory stimulation. *In vivo*, activity in SOM+ cells is state-dependent; for example, SOM+ cells are suppressed by VIP+ cells during whisking and locomotion (Fu et al. 2014; Gentet et al. 2012; Lee et al. 2013; Reimer et al. 2014). VIP+ cells also suppress SOM+ cells in auditory cortex in a state-dependent manner (Pi et al. 2013). Based on the results presented here, suppression of SOM+ cells by VIP+ cells in auditory cortex could act to prolong sound-induced network activity to facilitate integration of feedback input and sensory information from other modalities.

One potential caveat in interpreting the effects of suppressing SOM+ cells is that SOM+ cells strongly inhibit other interneuron types, including PV+ cells (Jiang et al. 2015; Pfeffer et al. 2013; Xu et al. 2013), and activation of SOM+ cells can have disinhibitory effects (Cottam et al. 2013; Xu et al. 2013). Therefore, we may underestimate the effects of SOM+ cell suppression on pyramidal cells because of disinhibition of PV+ cells and other interneurons, The importance of these types of network-level interactions has been highlighted recently in two reports. In the first (Seybold et al. 2015), activation of SOM+ and PV+ cells was shown to have diverse inhibitory effects on cells in the network depending on complex interactions between membrane properties at the singe cell level and network connectivity. A more recent study (Kato et al. 2017) showed SOM+ interneurons mediate a novel form of network-level lateral inhibition, and that inactivation of inhibitory cells could yield unexpected increases in inhibitory input to cells in the network. Further experiments involving targeted activation or suppression of subtypes of SOM+ cells may yield insights into these complex network-level effects. We note that the same caveat is less likely to arise for PV+ cells, as they mostly inhibit only other PV+ interneurons (Pfeffer et al. 2013), which would be simultaneously suppressed optogenetically in our experiments. Finally, we note another caveat as well: because spiking in interneurons contributes to measured population spiking activity, suppressing these cells will diminish modestly the observed effects on total population activity. Thus, although we observed increases in population activity with suppression of inhibitory cells (Figure 6), we may be underestimating those increases.

### Spike timing during network bursts

Timing information may exist not only relative to external stimuli but also relative to endogenous information packets or the phase of ongoing oscillations (Kayser et al. 2009; Luczak et al. 2013; Luczak et al. 2015). Our observation that pyramidal cells tend to fire at particular moments during network bursts (Figure 5) is consistent with these results. Suppression of PV+ cells led to earlier, briefer bursts of activity (Figure 6), and disrupted the organization of preferred firing phases of pyramidal cell during the burst (Figure 7). That is, with PV+ cell-mediated inhibition intact, activity within the population of pyramidal cells unfolded over a particular temporal sequence that spanned most of the duration of the network event; with PV+ cells suppressed, that organization was severely disrupted.

We used simulations to examine the consequences of this disrupted organization for decoding of temporal activity patterns. The tempotron model we employed was originally presented as a simple implementation of a logic unit able to capture the spatiotemporal input patterns characteristic of real nervous systems (Gütig and Sompolinsky 2006). However, such models cannot demonstrate how information is actually transmitted between networks in the brain, and using a single decoding unit oversimplifies a decoding process that is massively parallel at the output stage, as well as the input. Theoretical approaches using the tempotron model have allowed for estimates of how the encoding capacity of individual units varies with the spatiotemporal statistics of the input and highlights how timing information increases encoding potential (Rubin et al. 2010). We used an empirical implementation of the tempotron model to impose biologically relevant integration timescales on a hypothetical spatiotemporal discrimination task. We showed that when firing phases are concentrated at the beginning of bursts, as observed when PV+ cells are suppressed, different spike patterns are confused at a higher rate and take longer to learn (Figure 8), reducing the repertoire of distinct network responses.

### Effects on infragranular versus supragranular layers

Suppressing PV+ cells had greater effects on burst latency and burst peak in supragranular compared to infragranular layers (Figure 6). This observation suggests a role for these cells in regulating the flow of activity through the cortical column. Supragranular as well as infragranular pyramidal cells receive direct inputs from thalamus in auditory cortex (Ji et al. 2015; Krause et al. 2014), as in other sensory areas (Constantinople and Bruno 2013), but both spontaneous and stimulus-induced activity often originate infragranularly and then spreads to other layers (Beltramo et al. 2013; Chauvette et al. 2010; Krause et al. 2014; Stroh et al. 2013; Wester and Contreras 2012). Interestingly, when PV+ cells were suppressed, burst latency was nearly the same in supragranular and infragranular layers (Figure 6). This result supports a model in which PV+-mediated inhibition in the supragranular layers contributes to segregating the infragranular mechanisms generating network activity from supragranular information coding (Sakata and Harris 2009). In this model, sensory stimulation activates the network via dense-coding infragranular cells, with network activity spreading subsequently to other laminae with variable probability. Bursts restricted to the infragranular layers (Krause et al. 2014; Sakata and Harris 2009) could mediate motor responses in the absence of higher-order processing (Harris and Thiele 2011), while more complex and informative representations are revealed in the patterns of supragranular pyramidal cells that participate during induced network activity (Bathellier et al. 2012; Luczak et al. 2013). These data are consistent with suggestions that sparse coding in supragranular pyramidal cells (Barth and Poulet 2012; Crochet et al. 2011; DeWeese et al. 2003; Hromadka et al. 2008; Poulet and Petersen 2008; Sakata and Harris 2009) is likely regulated by inhibition (Bruno and Simons 2002; Crochet et al. 2011; Haider et al. 2010; Li et al. 2014). Consistent with this model, our simulations show that the effect of inhibition is greatest on spike patterns encoded by supragranular cells (Figure 9).

### Functional implications

Elucidating the roles of specific interneuron populations in regulating cortical network activity is central to understanding sensory information coding and its disruption under pathological conditions. Dysfunction of PV+-mediated inhibition is implicated in numerous conditions and disorders (Marín 2012), including schizophrenia (Lewis et al. 2012), autism (Hashemi et al. 2017), and epilepsy (DeFelipe 1999; Ogiwara et al. 2007). Consideration of spike timing with respect to informative packets of information can possibly resolve the ambiguity between rate coding of time-varying stimuli at a high temporal resolution versus true temporal codes (Borst and Theunissen 1999). Indeed, we show here that it is possible to preserve important timing relationships in isolated cortical slices, and specifically to dissociate temporal precision with respect to a stimulus and with respect to internal cortical activity. Our results suggest that PV+ interneurons have an important role in regulating the temporal structure of cortical network activity, and that this temporal structure should be considered as a potential target when inhibition is affected by behavioral state, attention, pharmacological interventions, or in cognitive disorders.

## Acknowledgements

The authors thank Sean Grady for technical support on this project and Harald Hentschke for comments on a draft of the manuscript.

---

**Supplementary Figure 1.**
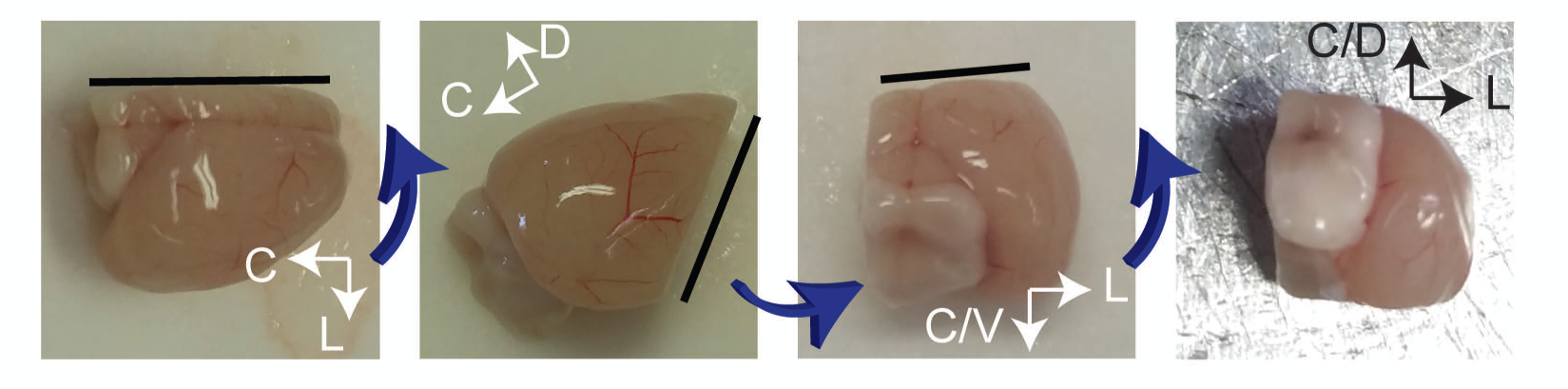

**Supplementary Figure 2.**
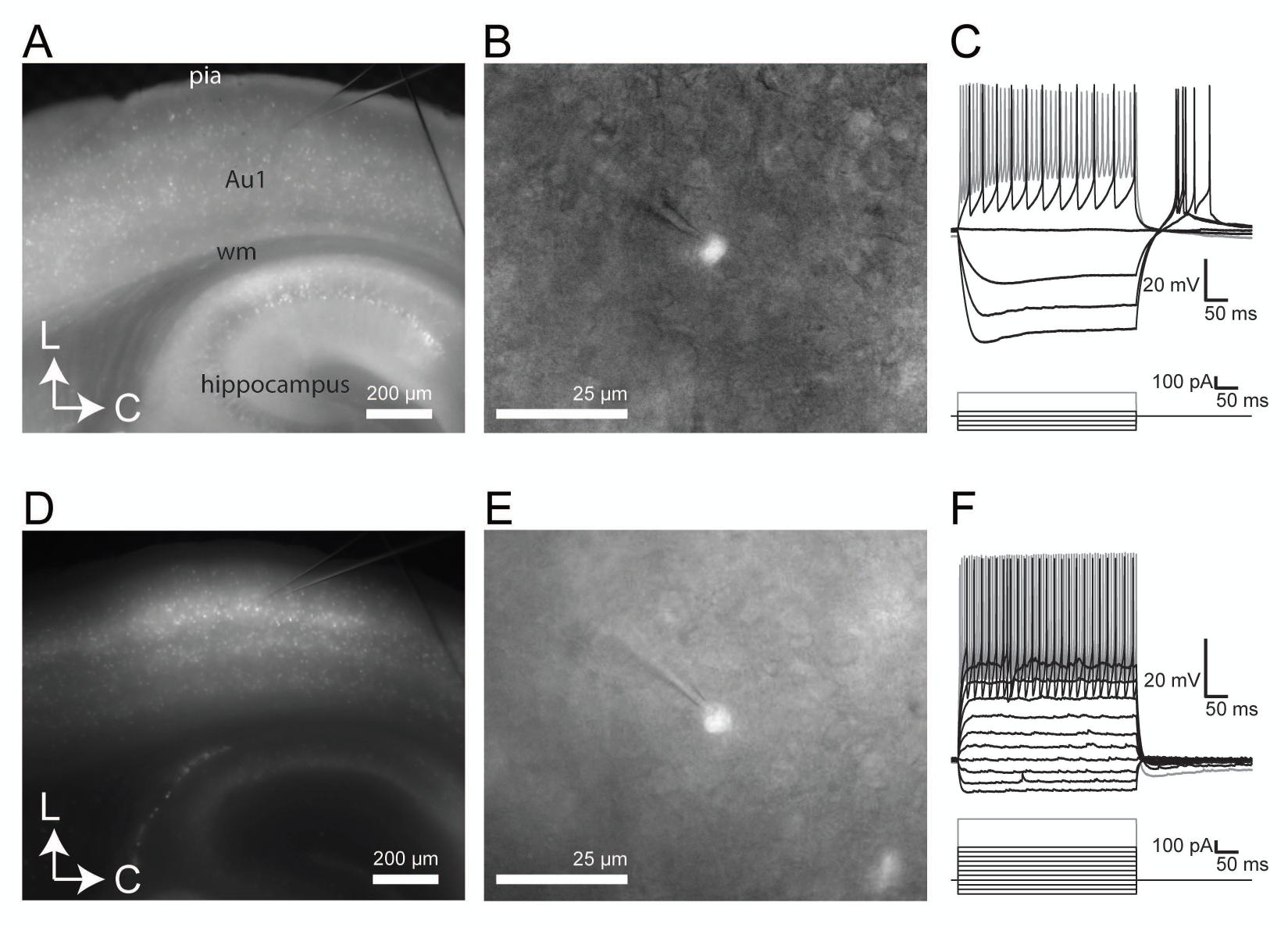

**Supplementary Figure 3.**
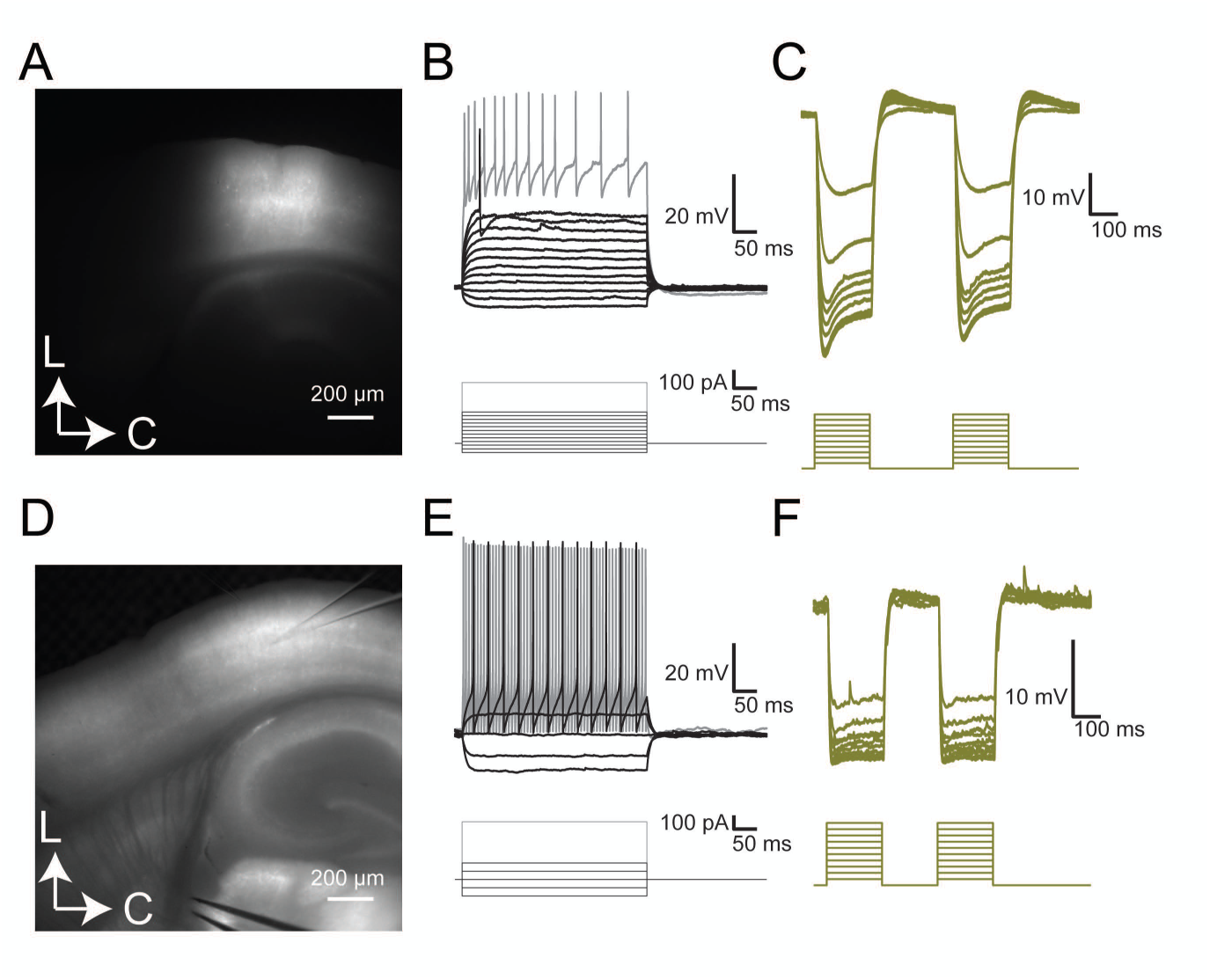

**Supplementary Figure 4.**
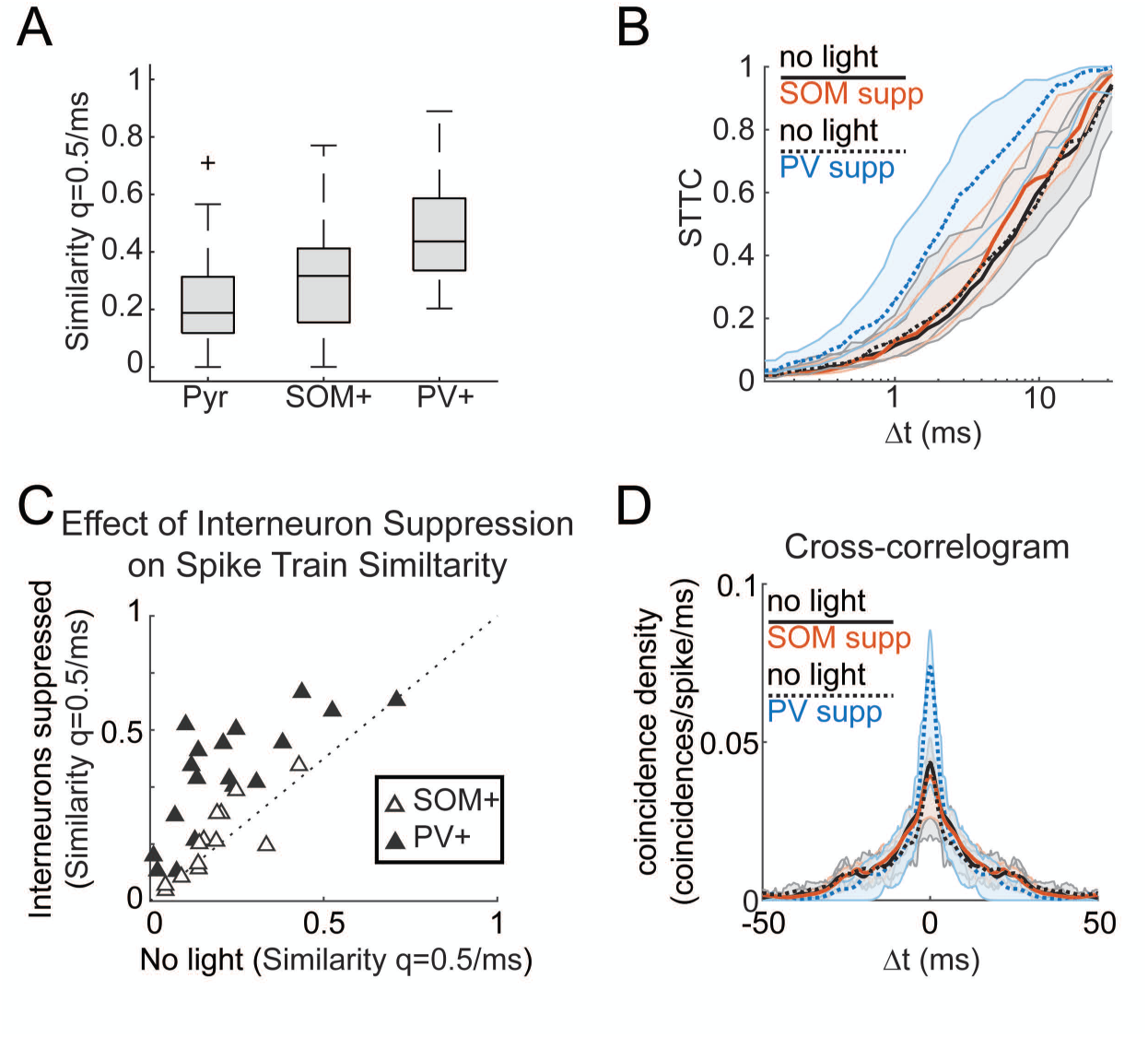

**Supplementary Figure 5.**
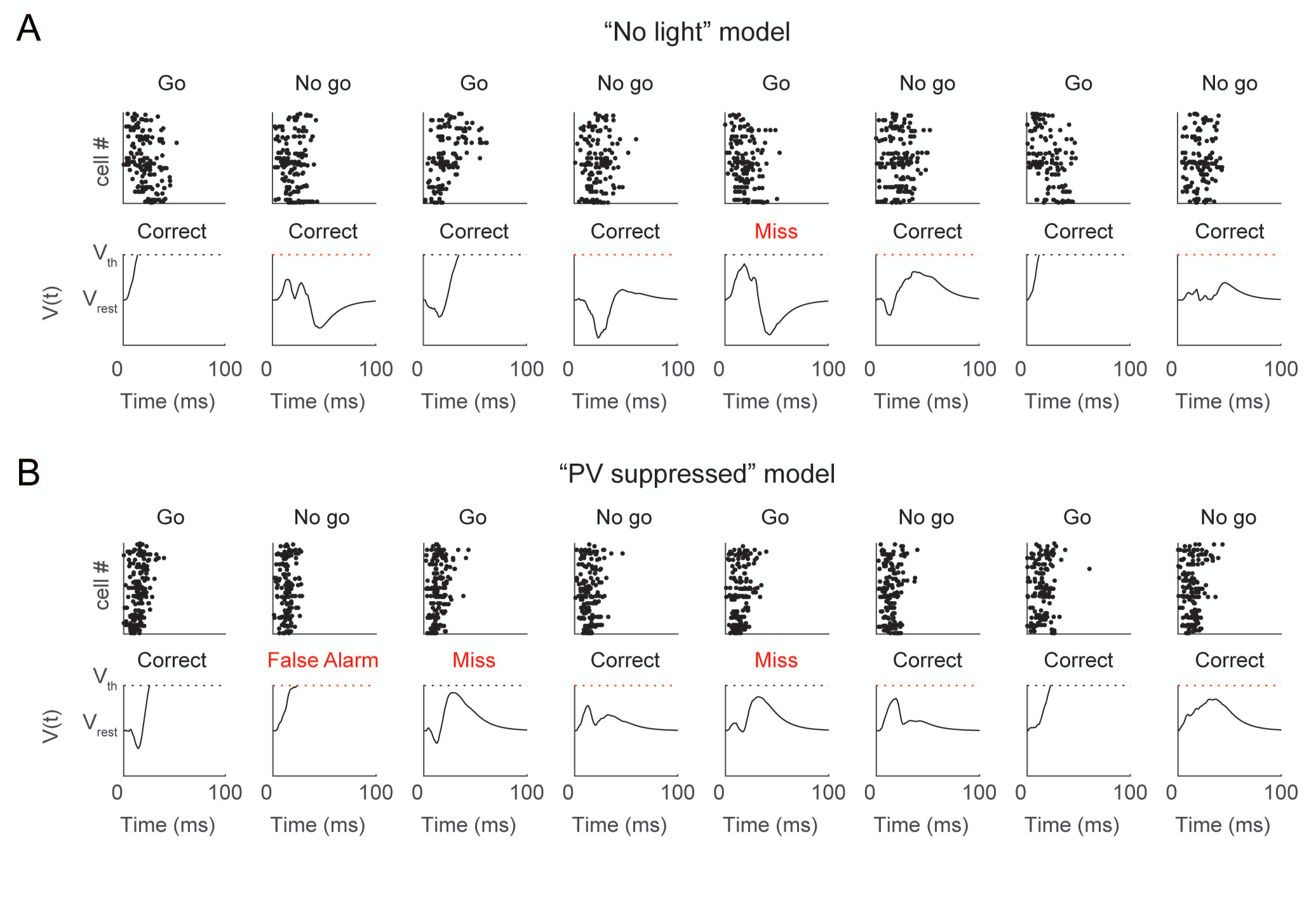

